# Autocrine Sfrp1 inhibits lung fibroblast invasion during transition to injury induced myofibroblasts

**DOI:** 10.1101/2022.07.11.499594

**Authors:** Christoph H. Mayr, Arunima Sengupta, Meshal Ansari, Jeanine C. Pestoni, Paulina Ogar, Ilias Angelidis, Andreas Liontos, Alberto Rodriguez-Castillo, Niklas J. Lang, Maximilian Strunz, Sara Asgharpour, Diana Porras-Gonzalez, Michael Gerckens, Bettina Oehrle, Valeria Viteri-Alvarez, Isis E. Fernandez, Michelle Tallquist, Martin Irmler, Johannes Beckers, Oliver Eickelberg, Gabriel Mircea Stoleriu, Jürgen Behr, Nikolaus Kneidinger, Ali Önder Yildirim, Katrin Ahlbrecht, Rory E. Morty, Christos Samakovlis, Fabian J. Theis, Gerald Burgstaller, Herbert B. Schiller

## Abstract

Fibroblast to myofibroblast conversion is a major driver of tissue remodeling in organ fibrosis. Several distinct lineages of fibroblasts support homeostatic tissue niche functions, yet, specific activation states and phenotypic trajectories of fibroblasts during injury and repair have remained unclear. Here, we combined spatial transcriptomics, longitudinal single-cell RNA-seq and genetic lineage tracing to study fibroblast fates during mouse lung regeneration. We discovered a transitional fibroblast state characterized by high Sfrp1 expression, derived from both Tcf21-Cre lineage positive and negative cells. Sfrp1+ cells appeared early after injury in peribronchiolar, adventitial and alveolar locations and preceded the emergence of myofibroblasts. We identified lineage specific paracrine signals and inferred converging transcriptional trajectories towards Sfrp1+ transitional fibroblasts and Cthrc1+ myofibroblasts. Tgfβ1 downregulated Sfrp1 in non-invasive transitional cells and induced their switch to an invasive Cthrc1+ myofibroblast identity. Finally, using loss of function studies we showed that autocrine Sfrp1 directly inhibits fibroblast invasion by regulating the RhoA pathway. In summary, our study reveals the convergence of spatially and transcriptionally distinct fibroblast lineages into transcriptionally uniform myofibroblasts and identifies Sfrp1 as an autocrine inhibitor of fibroblast invasion during early stages of fibrogenesis.

## Introduction

Extracellular matrix (ECM) producing myofibroblasts are an injury induced cell state that is a key therapeutic target to combat tissue fibrosis, one of the biggest unresolved clinical problems across most major chronic diseases. Single cell analysis has revealed substantial heterogeneity in fibroblast populations in multiple organs, which could be attributed to tissue and cell activation states, specific anatomical tissue niches, and lineage origin^1,2^. Several recent single cell RNA-seq studies described distinct subsets of collagen producing stromal cells in mouse and human lungs with distinct spatial locations and different functions in supporting epithelial repair^3–6^. It remains unclear though, how the lineage origin and spatial localization generate the functional heterogeneity of the scar producing myofibroblasts in pulmonary fibrosis.

The generation of fibrogenic myofibroblasts is generally associated with tissue repair and therefore also occurs transiently during normal lung regeneration^7^. However, in patients with non-resolving or even progressing interstitial lung disease, as seen in idiopathic pulmonary fibrosis (IPF), the ECM producing myofibroblasts persist and accumulate, thereby ultimately profoundly altering normal architecture and function of the lung. In addition to overproducing ECM matrices, the persistent myofibroblasts are thought to cause paracrine induction of ectopic aberrant epithelial cell states^8^, which may form a pathological cell circuit with the myofibroblasts^7,9–11^.

Genetic lineage tracing in mouse models provides evidence for an alveolar lipofibroblast to myofibroblast switch after lung injury that is reversible during the resolution of transient fibrosis upon completed epithelial regeneration^12,13^. If the alveolar lipofibroblast is an exclusive source of myofibroblasts in lung fibrosis, or if other cell types such as adventitial fibroblasts and pericytes also contribute, remains the subject of current investigations. Proliferation, evasion of apoptosis and invasive capacity of myofibroblasts are key hallmarks of fibrotic disease, and Tgfβ is a known master regulator of these processes^14,15^. A vast amount of literature demonstrates the Tgfβ induced decoration of contractile actomyosin bundles with α-smooth muscle actin (Acta2), the most widely used marker of mature myofibroblasts in tissues. However, fate mapping and immunofluorescence analysis of fibrotic tissues and myofibroblast foci also show that a substantial fraction of fibroblasts is Acta2 negative^16,17^, suggesting additional molecular complexity and heterogeneity amongst injury activated fibroblasts.

A recent single cell analysis of collagen producing cells in lung fibrosis revealed Cthrc1 as a specific marker of highly invasive Acta2+ myofibroblasts. These cells occur in both mouse and human lung and occupy fibroblastic foci in IPF^3^. Our study reveals the spatiotemporal evolution of distinct fibroblast states towards Cthrc1+ myofibroblasts, highlighting early events of injury induced fibroblast activation that precede myofibroblastic differentiation. We discover a novel Sfrp1+/Acta2− transitional state that is initially non-invasive and only becomes invasive upon Tgfβ driven differentiation towards the Cthrc1+ myofibroblast state. We show that Sfrp1 inhibits fibroblast invasion via induction of the RhoA pathway, which constitutes a novel pathway with potential for targeting fibrotic disease mediated by myofibroblasts.

## Results

### Heterogeneity of mesenchymal cells at distinct spatial localizations in the lung

The Tcf21+ lineage has been described to constitute the lung lipofibroblast population^18^. To study potential heterogeneity within the Tcf21+ and Tcf21− fibroblast lineages we tamoxifen labeled Tcf21+ cells in adult lungs at 11 weeks of age using Tcf21m-Crem-R26R-tdTomato mice (Suppl Fig. S1a). Tcf21 lineage positive cells were indeed found in the alveolar space in close proximity to AT2 cells and alveolar capillaries (Fig. S1b). We flow sorted Tcf21 lineage negative and lineage positive stromal cells (Epcam−/CD45−/CD31−/Lyve−) and performed single cell RNA-seq using the 10x genomics platform. After quality control filtering we recovered 12 068 cells from four individual mouse replicates and identified at least six distinct cell type identities with different lineage proportions (Fig. 1a and Suppl Fig. S1c-d). We used previously established marker genes^3,19,20^ to annotate the distinct clusters in our UMAP embedding (Markers in Table S1). All cell type identities were confirmed to express type 1 collagen (Col1a2) (Supl Fig. S2c) and were consistent with previous work^3^.

**Figure 1.**
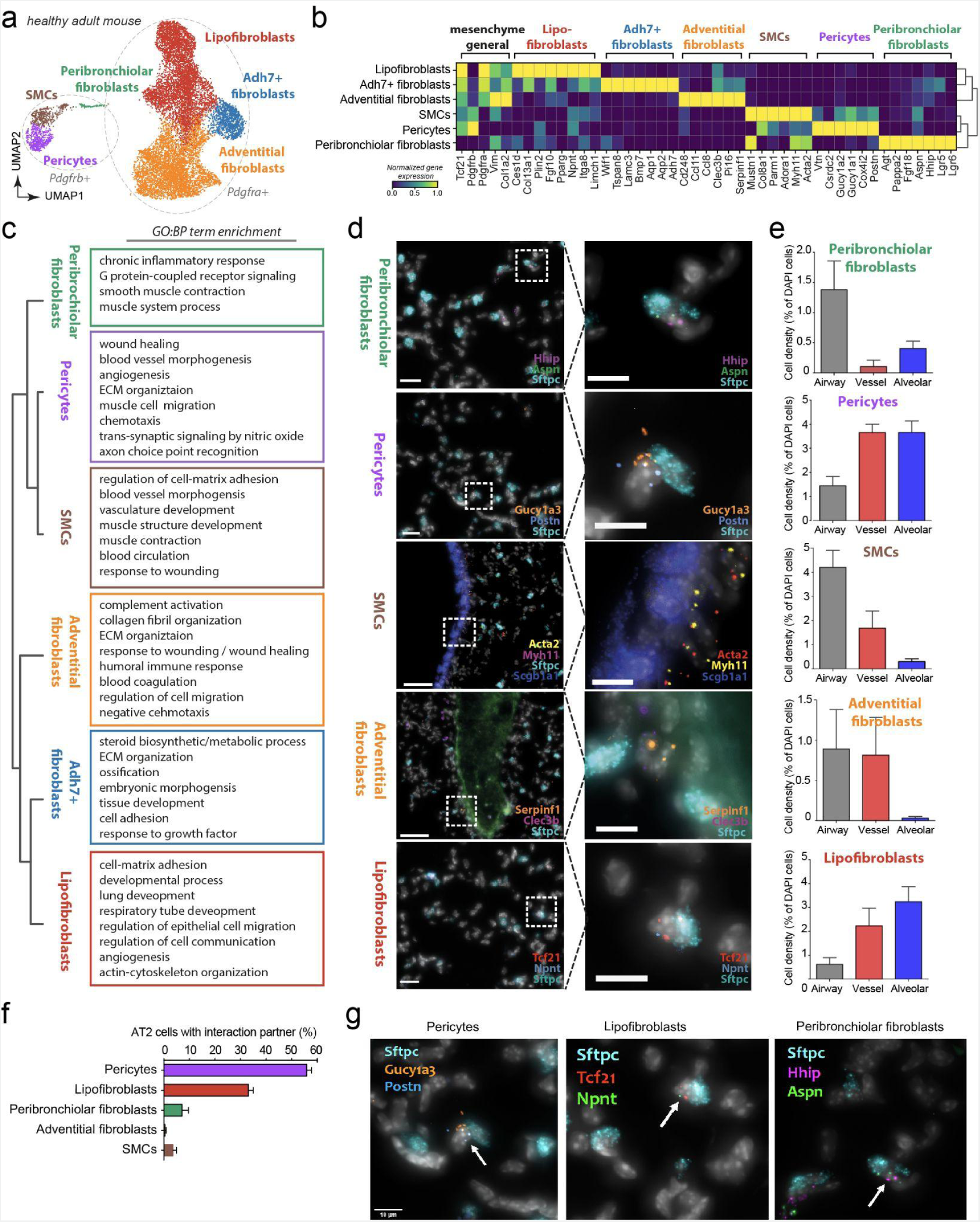
Stromal cell heterogeneity in the adult lung is associated with distinct spatial locations. **a)** UMAP clustering depicts the seven distinct mesenchymal cell types (n= 12068 cells) identified in the healthy adult mouse lung. Circles mark *Pdgfra* and *Pdgfrb* expressing cells. **b)** The seven mesenchymal cell types are characterized with distinct gene expression profiles, as depicted in the matrixplot. **c)** Marker genes of the indicated mesenchymal subtypes are enriched for characteristic GO:BP terms. FDR < 10%. Top 500 marker genes considered. **d)** Single mRNA multiplexed FISH (SCRINSHOT) with two cell type specific marker genes per cell type, as well as general markers allowed to identify preferential spatial locations of the mesenchymal cell types in the mouse lung. **e)** Localization analysis shows stereotypic localization of the mesenchymal cell types to peribronchiolar, adventitial and alveolar regions. The spatial location of 976 cells was quantified over 5 regions with 508 Pericytes, 70 Peribronchiolar cells, 206 Lipofibroblasts, 41 Adventitial fibroblasts and 151 smooth muscle cells (SMCs) counted. Graphs show cell density as percentage of cell type specific cells compared to all the cells (DAPI stain positive nuclei) in the area. Error bars represent SEM. **f)** Co-localization analysis of 1269 AT-2 cells revealed preferred interaction partners among the mesenchymal cell types. A total of 291 AT-2 interactions with mesenchymal cell types in the alveolar space were found: 167 Pericytes; 93 Lipofibroblasts, 15 Peribronchiolar fibroblasts; 3 Adventitial fibroblasts; 13 SMCs. **g)** SCRINSHOT pictures exemplifying the colocalization of AT-2 cells with distinct mesenchymal subtypes.

Within the Pdgfra+ cells (Fig. 1a) we identified lipofibroblasts expressing markers such as Plin-2 (Adrp), Col13a1, Npnt and high levels of Tcf21 (Fig. 1b). The marker genes of these lipofibroblasts were enriched for genes related to ‘lung development’, ‘cell-matrix adhesion’ and ‘angiogenesis’ (Fig. 1c). Adventitial fibroblast marker genes included Pi16, Clec3b and Serpinf1 (Fig. 1b) and were enriched for genes involved in ‘ECM organization’, ‘complement regulation’ and ‘wound healing’ (Fig. 1c). In addition, we identified a population of fibroblasts that featured expression of the All-trans-retinol dehydrogenase [NAD(+)] ADH7 and Bmp7, which we termed Adh7+ fibroblasts (Fig. 1b). The marker genes of these cells were interestingly enriched for ‘steroid metabolic processes’ (Fig. 1c), and a recent study localized such cells to adventitial cuffs^3^. Looking at Pdgfrb+ stromal cells we identified smooth muscle cells with Myh11 and Acta2 co-expression, as well as pericytes that expressed Gucy1a3 and Postn (Fig. 1b). Markers of both cell types were enriched for the term ‘blood vessel morphogenesis’ (Fig. 1c).

Finally, we also identified Pdgfra/Pdgfrb double positive cells (however with lower expression of both markers) as recently found also in mouse and human kidneys^21^. These cells expressed high levels of Hhip and Aspn and were enriched for ‘G-protein coupled receptor signaling’ as well as ‘smooth muscle contraction’ (Fig. 1b, c). The cells also specifically expressed Lgr5 and Lgr6 (Fig. 1b), which was previously reported to mark a peribronchiolar population of fibroblasts in the mouse^6^. Importantly, a recent study also identified a similar LGR5+ population of fibroblasts in distal human airways^22^. Sub-clustering analysis of this Hhip+ peribronchiolar fibroblast population revealed additional complexity with Lgr5/Lgr6 single as well as double positive populations (Supl Fig. S1e-g).

Notably, all cell types, with exception of the Hhip+ peribronchiolar fibroblasts, expressed Tcf21, however with highest expression in Lipofibroblasts, and were lineage labeled in the Tcf21m-Crem-R26R-tdTomato mouse (Supl Fig. S1c-d). This may indicate that Tcf21−lineage negative Hhip+ peribronchiolar fibroblasts constitute a developmentally distinct lineage from Tcf21+ stromal cells, and thus further work will be required to study the lineage history of Hhip+ peribronchiolar fibroblasts in lung development.

To associate the observed stromal cell heterogeneity with spatial location in the lung we designed a multiplexed mRNA FISH experiment using the recently developed SCRINSHOT method^23^. Using a set of common markers for epithelial cell types (e.g. Sftpc, Scgb1a1) as well as general stromal cell markers (e.g. Col1a2) as well as two specific markers for every stromal cell subtype we multiplexed the mRNA localization of 18 genes in 6 representative regions of adult murine lungs along the proximal distal axis of the airway tree (Fig. 1d and Supl Fig. 2a-c).

Quantification of 976 single cells that co-expressed the two marker genes (Fig. 1b, d), demonstrated a strong enrichment of Hhip+/Aspn+ peribronchiolar fibroblasts cells around airways, with some cells also showing an alveolar localization. Similarly, we found Myh11+/Acta+ smooth muscle cells and Serpinf1+/Clec3b+ adventitial fibroblasts enriched around airways and vessels. Gucy1a3+/Postn+ pericytes localized preferentially to the alveolar space and also around larger vessels. The Tcf21hi/Npnt+ lipofibroblasts finally were localized prefentially to alveolar space (Fig. 1e; Supl Fig. 2d-e). Consequently, the number of cells in close physical proximity (direct cell-cell contact) to Sftpc+ AT2 cells was highest for pericytes and lipofibroblasts with some Hhip+/Aspn+ cells also participating in the AT2 cell niche (Fig. 1f and Suppl. Fig. S2f). This highlights the complexity of the AT2 cell niche that we here demonstrate to contain at least 3 distinct stromal cell types.

### An activated fibroblast state characterized by high Sfrp1 and Col28a1 expression

To follow the fate of Tcf21−lineage positive and negative cells in lung fibrogenesis we flow sorted cells from Tcf21m-Crem-R26R-tdTomato mice (Suppl Fig. S1) at 14 days after bleomycin induced lung injury (Suppl Fig. S3 for gating strategy). In addition to the cell types observed in the ground state (Fig. 1), we identified three distinct clusters of injury induced myofibroblasts (Fig. 2a, b), all of which were a mixture of Tcf21−lineage positive and negative cells. Interestingly, we observed an expansion of Tcf21−lineage negative Hhip+ peribronchiolar fibroblasts (Supl Fig. S1j, p, q), and some of the markers of this population (e.g Lgr5 and Lgr6) were also expressed in a myofibroblast subset (Supl Fig. S1l), indicating that both Tcf21−lineage negative and positive cells converged into the myofibroblast phenotypes.

**Figure 2.**
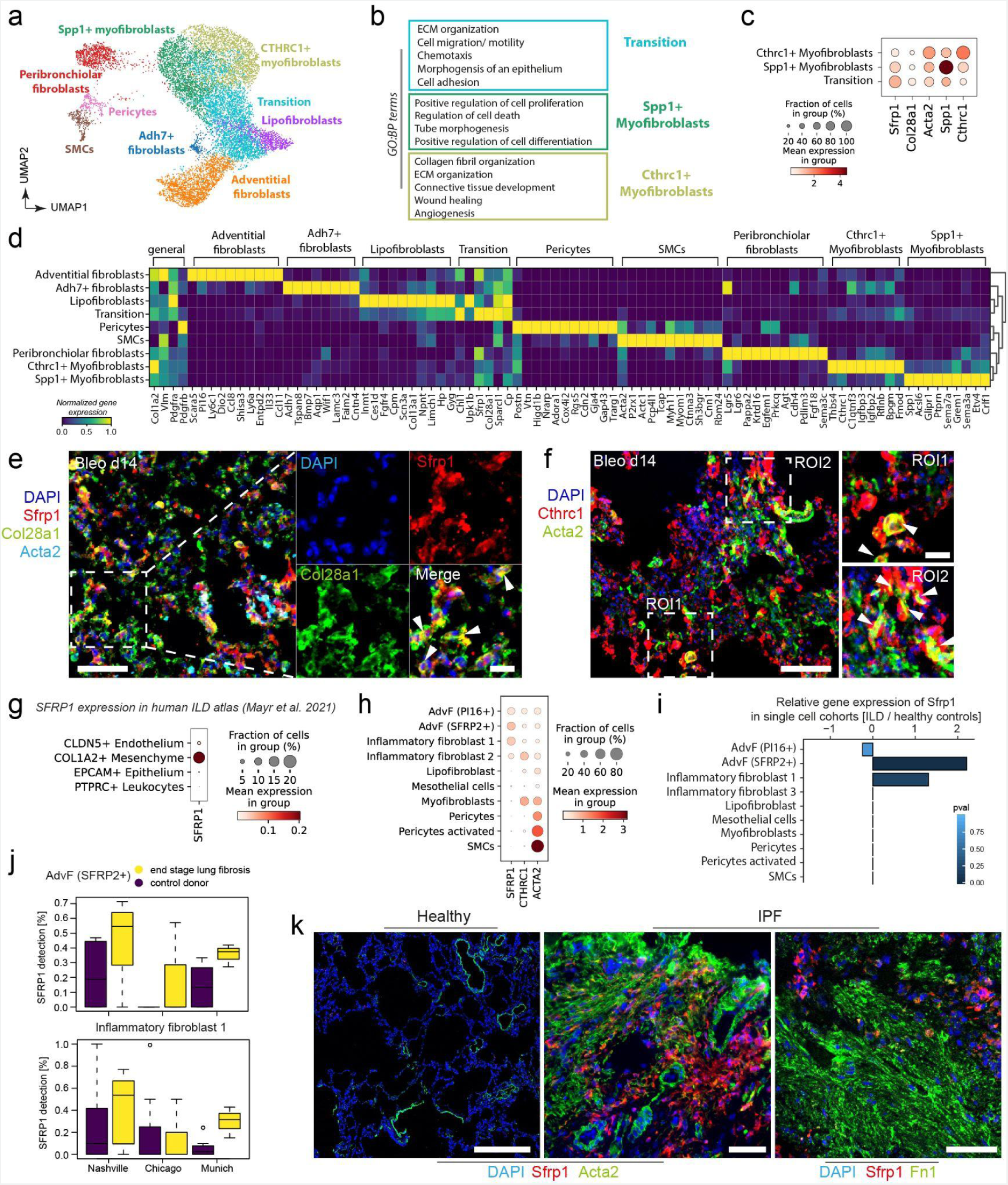
Three distinct activated mesenchymal cell types emerge after bleomycin induced mouse lung injury. **a)** UMAP clustering depicts mesenchymal cell types (n=12254 cells) identified at day 14 after bleomycin injury. **b)** The marker genes of activated mesenchymal subtypes are enriched for characteristic GO:BP terms. FDR < 10%. Top 500 marker genes considered. **c)** The dotplot depicts a gradient of marker gene expression across the activated cell types. **d)** The mesenchymal cell types are characterized with distinct gene expression profiles, as depicted in the matrixplot. **e-f)** Immunofluorescence analysis of lung tissue sections from bleomycin-treated mice demonstrate colocalization of upregulated **e)** Sfrp1(red) and Col28a1 (green) fourteen days (Bleo d14) after injury. Arrowheads in the magnified insets indicate Sfrp1/Col28a1 double positive cells. Sfrp1/Col28a1 double-positive cells did not co-localize with Acta2 (cyan). Scale bars = 100 µm and 50 µm. **f)** At d14 after injury, Acta2 (green) and Cthrc1 (red) double-positive cells were found (indicated by arrowheads in the magnified insets for ROI1 and ROI2 in the very right panel). Nuclei in blue color (DAPI). Scale bars = 100 µm and 20 µm. **g-j)** Distinct markers from the bleomycin mouse model were analyzed in a recently published integrated atlas of human lung fibrosis at single cell resolution^24^. **g)** Sfrp1 showed specific expression on human Col1a2+ positive mesenchymal cells. **h)** Marker gene expression across all mesenchymal cell types identified in the human lung. **i-j)** Differential gene expression analysis between ILD patient samples and healthy control samples, **(i)** reveals significant regulation of Sfrp1 in Adventitial fibroblasts and Inflammatory fibroblasts 2. **j)** This regulation is consistent across all three cohorts in the atlas. **k)** Immunofluorescence analysis of lung tissue sections from healthy controls and IPF patients demonstrates no colocalization of Sfrp1 with myofibroblasts markers Acta2 or fibronectin (Fn1). Scale bars 200 µm, 100 µm and 100 µm, respectively.

As previously reported^3^, we identified a cluster of Cthrc1+/Acta2+ myofibroblasts that was enriched for terms such as collagen fibril organization and wound healing (Fig. 2b). We also noted a quite distinct cluster of Acta2+ myofibroblasts that featured enhanced expression of Spp1 (Fig. 2 c,d). In addition, one cluster featured co-expression of myofibroblast genes with markers of different fibroblast types in the uninjured lung, most notably the lipofibroblasts, and we therefore termed these cells transitional fibroblasts, which featured many genes involved in regulation of cell locomotion (Fig. 2b,c,d). Several marker genes, including the secreted frizzled-related protein 1 (Sfrp1) and the collagen type 28 (Col28a1), showed highest expression in this putative intermediate cell population (Fig. 2c,d; markers in Table S2).

The Sfrp1+ transitional fibroblasts showed lower expression of Acta2 than the Cthrc1+ and Spp1+ fibroblasts. Of note, using whole tissue proteomics by mass spectrometry we have previously identified Col28a1 and Sfrp1 as a transiently induced protein after bleomycin injury^25^ (Suppl Fig. S4c). Gene-gene correlation of Sfrp1 with other genes across all single cells and time-points revealed a core gene set, including Col28a1, associated with the Sfrp1+ transitional fibroblasts (Suppl Fig. S4d). Co-staining Sfrp1 and Col28a1 confirmed co-expression in Acta2 low or negative transitional cells (Fig. 2e), while Cthrc1 positive cells expressed higher levels of Acta2 (Fig. 2f).

We analyzed cross species conservation of cell type identities in human lungs and compared Col1a2+ cells after bleomycin injury with COL1A2+ cells in ILD patients from three independent single cell studies on pulmonary fibrosis^24^. Sfrp1 was specific to Col1a2+ mesenchymal cells and was only weakly expressed in some Cldn5+ endothelial cells (Fig. 2g). Expression was restricted to adventitial fibroblasts as well as disease enriched “inflammatory” subsets of fibroblasts with high expression of cytokines and chemokines (Fig. 2g, h). The increased expression of Sfrp1 in these disease induced fibroblast states was independently present in all three study cohorts (Fig. 2i,j). Finally, immunostainings confirmed increased expression of SFRP1 in ACTA2 and FN1 negative cells in the IPF patients (Fig. 2k).

### Sfrp1+ transitional fibroblasts precede the appearance of Cthrc1+ myofibroblasts

To follow the transcriptional dynamics of all Col1a2+ stromal cells during the inflammatory, fibrogenic and resolution phase of lung regeneration we collected Epcam−/Pecam1−/Lyve1−/CD45− stromal cells at 18 timepoints after bleomycin induced lung injury (Fig. 3a-c). This longitudinal single cell RNA-seq dataset of Col1a2+ cells (n= 69185 cells) at 18 different timepoints after injury (Fig. 3b), again featured three distinct injury induced cell states with similar marker genes (Fig. S4a) as observed in the Tcf21−lineage tracing dataset above (Fig. 3c; markers in Table S3).

**Figure 3.**
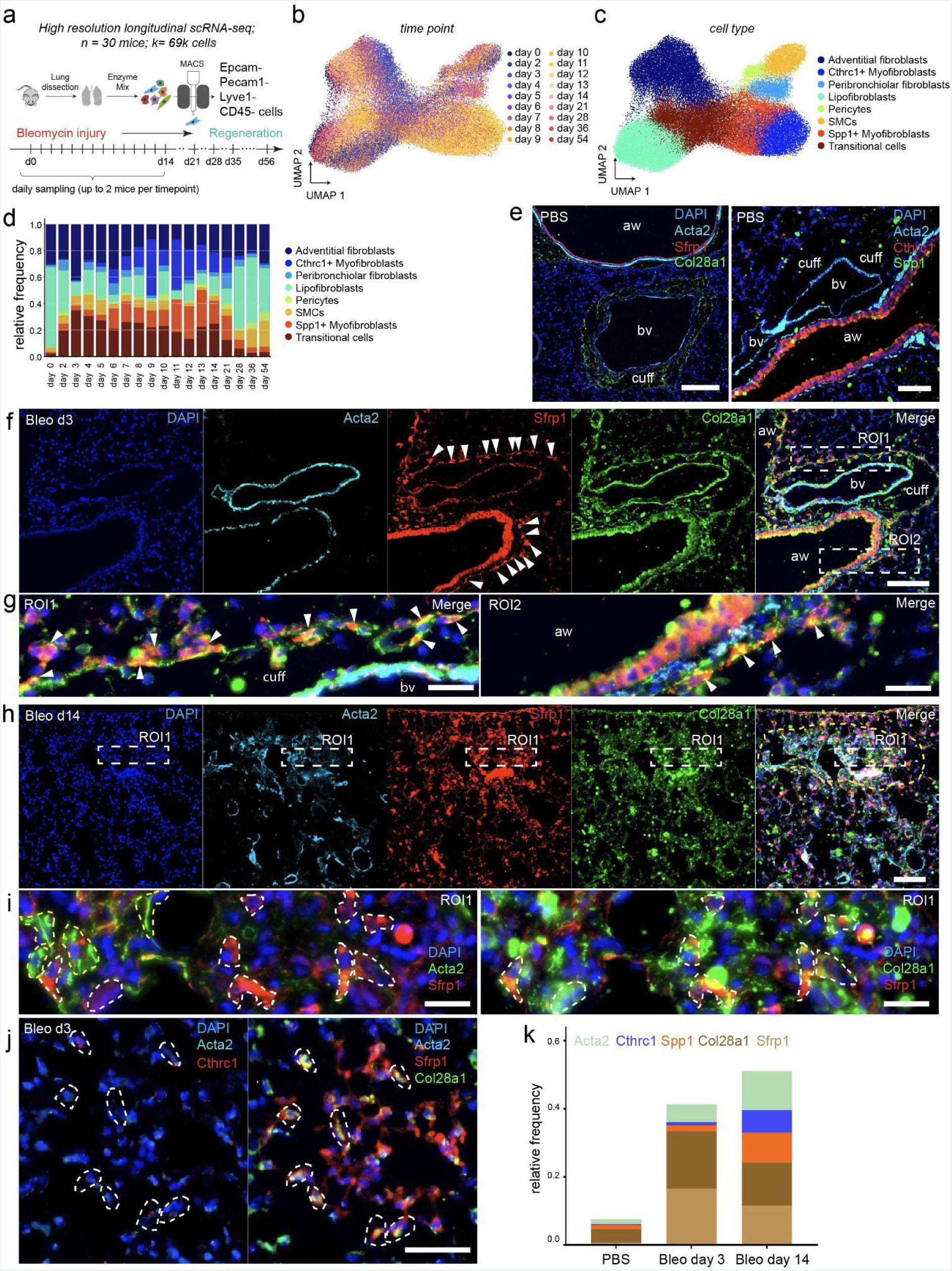
Sfrp1+ transitional fibroblasts emerge in adventitial and alveolar space upon injury preceding the emergence of Cthrc1+ myofibroblasts. **a)** A high resolution longitudinal data set was generated by subjecting MACS sorted cells from the mesenchymal compartment to scRNAseq at the 18 indicated time points. UMAP embedding displays cells colored by **b)** time point and **c)** cell type identity. **d)** The bar blot shows the distribution of cell type frequencies across time points. **e)** Immunofluorescence analysis of lung tissue sections from PBS control mice. In the left panel cell nuclei were immunostained with DAPI (blue) and SMCs with Acta2 (cyan.) Sfrp1 (red) staining was absent apart from nonspecific signals in bronchial epithelial cells. Col28a1 (green) was mostly found as a filamentous staining in “cuffs” surrounding blood vessels (bv). In the right panel cell nuclei were stained with DAPI (blue) and SMCs with Acta2 (cyan). Cthrc1 (red) staining was absent apart from nonspecific signals in bronchial epithelial cells. Spp1 is depicted in green. aw = airways and bv = blood vessels. Scale bars = 100 µm. **(f)** Immunofluorescence analysis of lung tissue sections from bleomycin-treated mice at day 3 after injury (Bleo d3) demonstrating appearance of Sfrp1 (red) positive cells surrounding blood vessels (bv) and airways (aw) (arrowheads). Scale bar = 100 µm. **(g)** Arrowheads in the magnified insets taken from ROI1 and ROI2 of Fig.4f indicate Sfrp1/Col28a1 double positive cells surrounding blood vessels (bv) in ROI1 reminiscent of adventitial fibroblasts, was well as surrounding airways (aw) in ROI2 reminiscent of peribronchiolar fibroblasts. Scale bars = 20 µm. **(h)** Yellow dashed-line indicates a fibrotic region of lung tissue after 14 days of bleomycin treatment. Acta2 staining in cyan exhibits the appearance of myofibroblasts, and concomitant expression of Sfrp1 (red) and Col28a1 (green) in this fibrotic region. Cell nuclei stained with DAPI in blue. Scale bar = 100 µm. A magnified view of fibrotic ROI1 is displayed in **(i)**, in the left panel demonstrating mutually exclusive appearance of Sfrp1+ (red and dashed white lines) and Acta2+ (green and dashed yellow lines) cells. Here, for better detection of colocalization signals between Acta2 and Sfrp1, we switched colors of Acta2 from cyan as depicted in (h) to green. The right panel denotes Sfrp1 (red) and Col28a1 (green) double positive cells encircled with white dashed lines. Cell nuclei stained with DAPI in blue. Scale bars = 20 µm. **(j)** Iterative immunofluorescence staining of parenchymal lung tissue (4i-FFPE) at day 3 after injury indicating Col28a1 (green)/Sfrp1 (red) double positive cells in the left image. The same cells, as indicated by white dashed lines, were found to be Cthrc1 (red) and Acta2 (cyan) negative. Scale bar = 50 µm. A larger overview image can be found in Supl. Fig. S7a. **(k)** Stacked bar-plot denoting relative frequencies of Acta2, Cthrc1, Spp1, Col28a1 and Sfrp1 positive cells at day 3 and day 14 after bleomycin treatment and compared to PBS controls from software-based segmented images as exemplified in Supl. Fig. S7c. Three different ROIs (each 1.1 mm^2^ in size) from two different mice for each condition were analyzed.

The relative frequency of Sfrp1+ transitional fibroblasts peaked early in the time-course at day 3 after injury, preceding the increase in frequency of Spp1+ and Cthrc1+ myofibroblasts at later timepoints with their peak around day 9 to day 21 (Fig. 3d). The appearance of Sfrp1+ transitional cells was inversely correlated with a reduction in the baseline states (e.g. lipofibroblasts and adventitial fibroblasts). Importantly, this dynamic was reverted back to baseline from day 28 onwards and we observed a reduction of the myofibroblast states and concomitant increase of the baseline states (Fig. 3d). Proliferation analysis (Mki67/Top2a co-expression) showed a transient increase in proliferation rates, with most of the proliferating cells within the Spp1+ and Cthrc1+ myofibroblasts, which was reverted back to baseline from day 28 onwards (Suppl Fig. S4b). The proliferation analysis indicated that the increase of Sfrp1+ cells early after injury was not due to expansion of a pre-existing pool of these cells but rather the dedifferentiation of baseline fibroblast states.

We next validated the early appearance of Sfrp1+ transitional fibroblasts preceding Spp1+ and Cthrc1+ myofibroblasts during the time-course of injury and repair using immunofluorescence analysis (Fig. 3e-k). Signals for Sfrp1, Cthrc1 and Spp1 were mostly absent in healthy controls, except for nonspecific signals marking luminal areas of the airway epithelium (Fig. 3e). In healthy lungs, Col28a1 was detected as filaments around bronchovascular cuffs, whereas Acta2 was primarily observed in smooth-muscle cells next to airways (aw) and blood-vessels (bv) (Fig. 3e). The localization of adventitial and peribronchial fibroblasts within and next to bronchovascular cuffs was described before^3^ and is in agreement with our smFISH analysis (Fig. 1).

On day three post injury, we observed Sfrp1 and Col28a1 double-positive transitional cells emerging around airways and blood-vessels, while Acta2 was still mainly expressed in bronchovascular smooth-muscle cells (Fig. 3f and Fig. 3g). Similarly, we observed Sfrp1+ transitional cells appearing early after injury in alveolar areas devoid of Cthrc1+ myofibroblasts (Fig. 3j and Supl. Fig. S7a,b). In contrast, at day 14 post injury, we detected both Acta2 positive myofibroblasts as well as cells with high Sfrp1 and Col28a1 expression (Fig. 3h, i). At day 3 post injury we observed mainly Sfrp1 and Col28a1 double-positive transitional cells, which were clearly negative for the myofibroblast markers, as evidenced by multiplexed immunofluorescence using iterative stainings (4i-FFPE) (Fig. 3j). The relative immunofluorescence signal of Spp1 and Cthrc1 was clearly increased in fibrotic regions at day 14 after injury (Supl. Fig. S7c). A quantitative analysis of 1.1 mm^2^ large tissue areas by software-based segmentation of immunofluorescence images confirmed the rise of Sfrp1+ and Col28a1+ double-positive transitional cells at day 3, preceding the expansion of Spp1+ and Cthrc1+ myofibroblasts until day 14 after injury (Fig. 3k). Of note, our time-resolved assessment of cell type proportions by imaging explicitly confirmed the cell type frequencies in the single cell expression data as presented in Fig. 3d.

### Convergence of multiple mesenchymal cell types towards myofibroblast identity

To infer lineage relationships between the mesenchymal cell types and activation states we used CellRank^26^ to analyze the longitudinal injury dataset with 18 distinct timepoints after injury (Fig. 4). CellRank combines trajectory inference algorithms with directional information from RNA velocity vectors overcoming limitations of simple 2D projections of local RNA velocity vectors.

**Figure 4.**
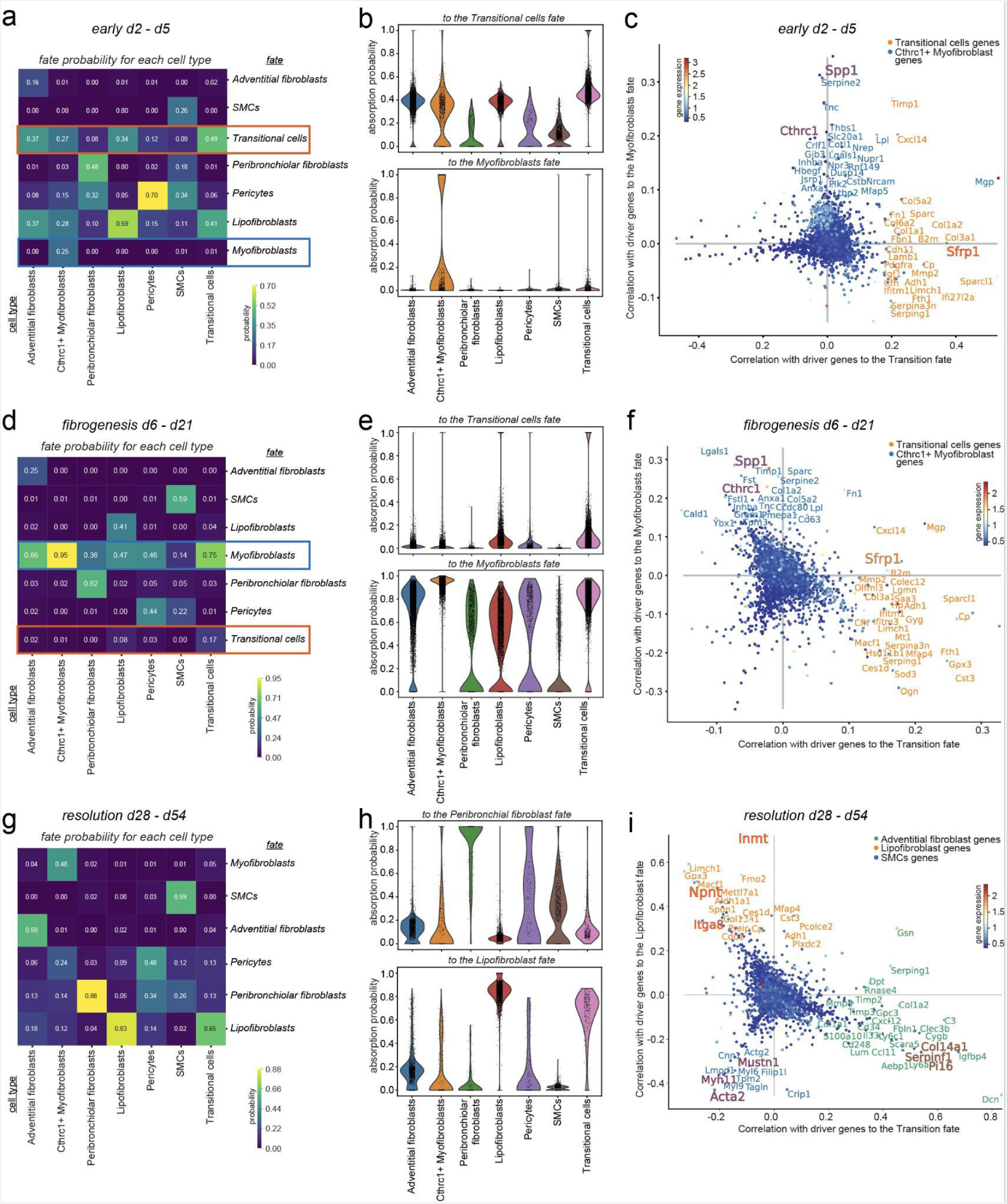
Lineage convergence and two-way conversion of fibroblasts in lung regeneration. **a,d,g)** CellRank’s coarse-grained and directed transition matrix was calculated for the given three phases of the bleomycin lung injury time course, of early phase (day 2 to day 5), fibrogenesis phase (day 6 to day 21) and resolution phase (day 28 to day 54). Terminal macrostates were set manually and annotated according to their overlap with the underlying gene expression clusters. The heatmap represents the mean absorption probabilities of every cell in the given cell types to the terminal fates. **b,e,h)** Violin plots indicating the adsorption probability towards the specified fate for every cell in all the cell types. **c,f,i)** Scatter plots show top driver genes, predicted to facilitate transition to the given terminal fate, visualized as gene-correlations between two terminal fate lineages. The top genes of a third lineage are visible by the third color of the gene names. Dots are colored according to their mean gene expression.

We separated the bleomycin time course data in the three phases of early injury (Fig. 4a,b,c), fibrogenesis (Fig. 4d,e,f) and resolution (Fig. 4g,h,i), and computed CellRank’s directed transition matrices separately. The identified macrostates were set as terminal states, so that every cell type was represented with a fate, corresponding to its ground state identity. We next computed fate probabilities for every cell and grouped them by cell types, to show their specific potential to differentiate towards the terminal states. The results were summarized in heat maps showing the fate probability for each cell type (Fig. 4a,d,g) and in violin plots for selected fates, visualizing the probability distribution among the cells in each cell type (Fig. 4b,e,h). Finally, we used again CellRank to predict trends for key lineage driver genes, that show upregulated expression along the lineage trajectory towards the terminal states and compared the genes along different trajectories using scatter plots (Fig. 4c,f,i).

Analysis of fate probabilities in the early phase of injury (d2-d5) showed that adventitial fibroblasts and lipofibroblasts had a comparably high fate probability towards the Sfrp1+ transitional state, while all other cell types only showed high probabilities to their own identity (Fig. 4a,b). Thus, this analysis indicates that after injury the adventitial fibroblasts and lipofibroblasts have the highest propensity to differentiate towards the Sfrp1+ transition state, rather than staying in their ground state. Probabilities towards the myofibroblast were still low and only represented by the few Cthrc1+ myofibroblast already present in this early injury phase (Fig. 4a,b). Top driver genes towards transitional cells were characterized by various collagens, chemokines and notably Sfrp1 as one of the most highly correlating driver genes, while the myofibroblast lineage showed a mix of injury marker genes including Tnc, Spp1 and Thbs1 (Fig. 4c).

In the fibrogenesis phase (d6-d21), the overall fate probabilities had shifted away from the transition state to the myofibroblast fate, with Sfrp1+ transitional cells showing the highest probability among non-Cthrc1+myofibroblast cells, suggesting a differentiation of the transitional cells into myofibroblasts (Fig. 4d,e). The general rise in fate probabilities towards myofibroblasts amongst the Pdgfrb+ mesenchymal cell population could also be attributed to the increase in frequency of myofibroblasts during that phase. Top driver genes towards myofibroblasts represented classical myofibroblast-associated genes including ECM proteins Lgals1, Sparc, and Spp1 as well as Cthrc1 (Fig. 4f).

In the resolution phase, CellRank predicted high probabilities of Cthrc1+ myofibroblast differentiation towards lipofibroblast, Hhip+ peribronchiolar fibroblast and pericyte states, while the remaining Sfrp1+ transitional cells had the highest probability towards lipofibroblasts (Fig. 4g). This prediction is in line with previous observations of a two-way conversion between lipogenic and myogenic fibroblastic phenotypes^12^. All other cell types were predicted to keep their identity (Fig. 4h). Therefore, the lineage driver genes in this resolution phase were characterized by the respective baseline cell state marker genes for each cell type (Fig. 4i).

To analyze gene expression dynamics over the entire time course we used a spline regression model that revealed genes with differential expression along the time-course in at least one cell type (Table S4). We found 25 different collagens with dynamic expression patterns after injury or between the different stromal cell types (Fig. 5a). As previously reported Col13a1 and Col14a1 were found to be very specific for lipofibroblasts and adventitial fibroblasts respectively. Col18a1 was specific to pericytes, Col5a3 specifically marked Cthrc1+ myofibroblasts, and Col28a1 marked the Sfrp1+ transitional as well as less prominently the Spp1+ myofibroblasts (Fig. 5a).

**Figure 5.**
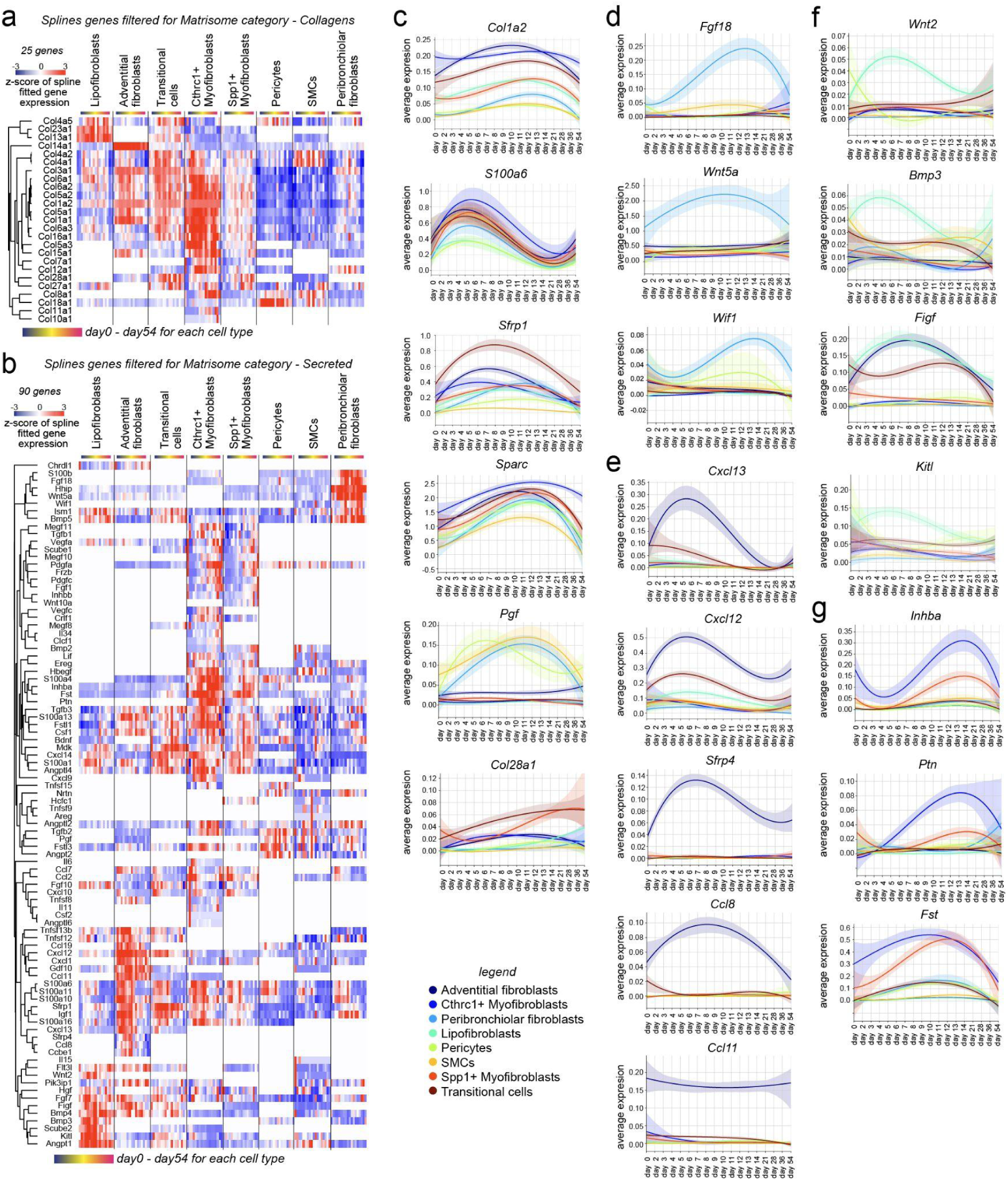
Highly specific expression changes in distinct fibroblast lineages. **a-b)** Heat maps display genes that showed differential expression along the time-course in at least one cell type using a spline regression model for **a)** 25 different collagens and **b)** 90 secreted matrisome^27^ genes. **c-g)** Line plots show average expression of indicated genes for each cell type along the bleomycin time course, for **c)** general mesenchymal marker genes, **d)** genes regulated over time in peribronchial fibroblasts, **e)** adventitial fibroblast, **f)** lipofibroblasts, and **g)** Cthrc 1+ mMyofibroblasts.

We also identified 90 secreted matrisome^27^ genes with significant expression changes along the injury time-course (Fig. 5b). All cell types transiently increased their expression of Collagen type-I (Col1a2), even though from very different baseline expression levels (Fig. 2f, h). Similarly the ECM protein Sparc, which is typically used as a marker for myofibroblasts^28^, showed a strong transient upregulation in all mesenchymal cell types (Fig. 5c). We also found common early events in fibroblast activation such as the upregulation of S100a6 early after injury (Fig. 5c). Other factors were specific to groups of cell types, like the placenta growth factor (Pgf), which was upregulated upon injury exclusively in smooth muscle cells, pericytes, and Hhip+ peribronchiolar fibroblasts, indicating a function in supporting the specific peribronchiolar vascular niche. Pgf binds the receptor Flt1/Vegfr-1, indicating an instructive role for endothelial cells in the repair process.

Peribronchiolar fibroblasts specifically featured the secretion of important morphogens such as the Hedgehog interacting protein (Hhip), the Wnt inhibitory factor 1 (Wif1), the fibroblast growth factor 18 (Fgf18), and Wnt5a (Fig. 5d). Fibroblast derived Wnt-signaling has been shown to define a specific AT2 cell niche in normal homeostasis, with a key role of Wnt5a in this fibroblast niche^29^. Our data demonstrates that stromal cell Wnt5a is exclusively derived from the Tcf21−Cre lineage negative Hhip+ fibroblasts with transiently increased expression after injury (Fig. 5d). WNT5A was also part of a distinct mesenchymal niche in human distal airways and secreted by LGR5+ fibroblasts^22^. Our smFISH analysis showed that a subset of Hhip+ peribronchiolar fibroblasts also localized to alveolar space in close proximity to AT2 cells (Fig. 1e, f), suggesting additional heterogeneity in this lineage of cells. Importantly, Hhip+ peribronchiolar fibroblasts also showed a transient upregulation of the transitional cell type marker Sfrp1 over time, indicating activation and potential to transition to myofibroblasts (Supl Fig. S5).

Adventitial fibroblasts were a specific source of the the Wnt modulator Sfrp4, as well as chemokines induced by injury, such as Cxcl13 to recruit B-cells, and Cxcl12 and Ccl8 to recruit T-cells, monocytes and neutrophils. Interestingly, the chemokine Eotaxin (Ccl11) was a specific marker gene for adventitial fibroblasts that remained unaltered by injury (Fig. 5e). Lipofibroblasts specifically increased the expression of the important morphogens Wnt2 and Bmp3, as well as the stem cell factor SCF (Kitl) (Fig. 5f). The SCF-c-Kit pathway has been shown to be activated in bleomycin-injured lungs, with potential profibrotic effects via recruitment of Kit+ immune cells to the lung^30^.

In summary, we found Sfrp1+ transitional cells in peribronchiolar, adventitial and alveolar regions already early after injury, preceding the emergence of Cthrc1+ myofibroblasts. Different mesenchymal cell types showed highly distinct patterns of morphogen, chemokine/cytokine, and ECM expression patterns after injury, indicating specific functions in the ensuing repair process. Trajectory inference analysis within our high resolution longitudinal dataset suggests convergence of several mesenchymal cell types towards Sfrp1+ transitional cells, which ultimately are predicted to give rise to Cthrc1+ myofibroblasts. Furthermore, our analysis indicates a reverse conversion of Cthrc1+ myofibroblasts towards lipofibroblasts, as well as possibly peribronchiolar fibroblasts and pericytes in the resolution phase upon completed epithelial regeneration.

### TGF-beta mediates differentiation of Sfrp1+ transitional fibroblasts into myofibroblasts

To infer potential regulators of the fibroblast cell state transitions after injury we used NicheNet^31^, which links ligands in cell-cell communication networks to changes in gene expression in receiving cells based on prior knowledge on downstream target genes in cells. NicheNet predicted the ligands with highest probability to induce expression of the top driver genes from the CellRank outputs (Fig. 5) for Sfrp1+ transitional cells and Cthrc1+ myofibroblasts respectively (Fig. 6a, b).

**Figure 6.**
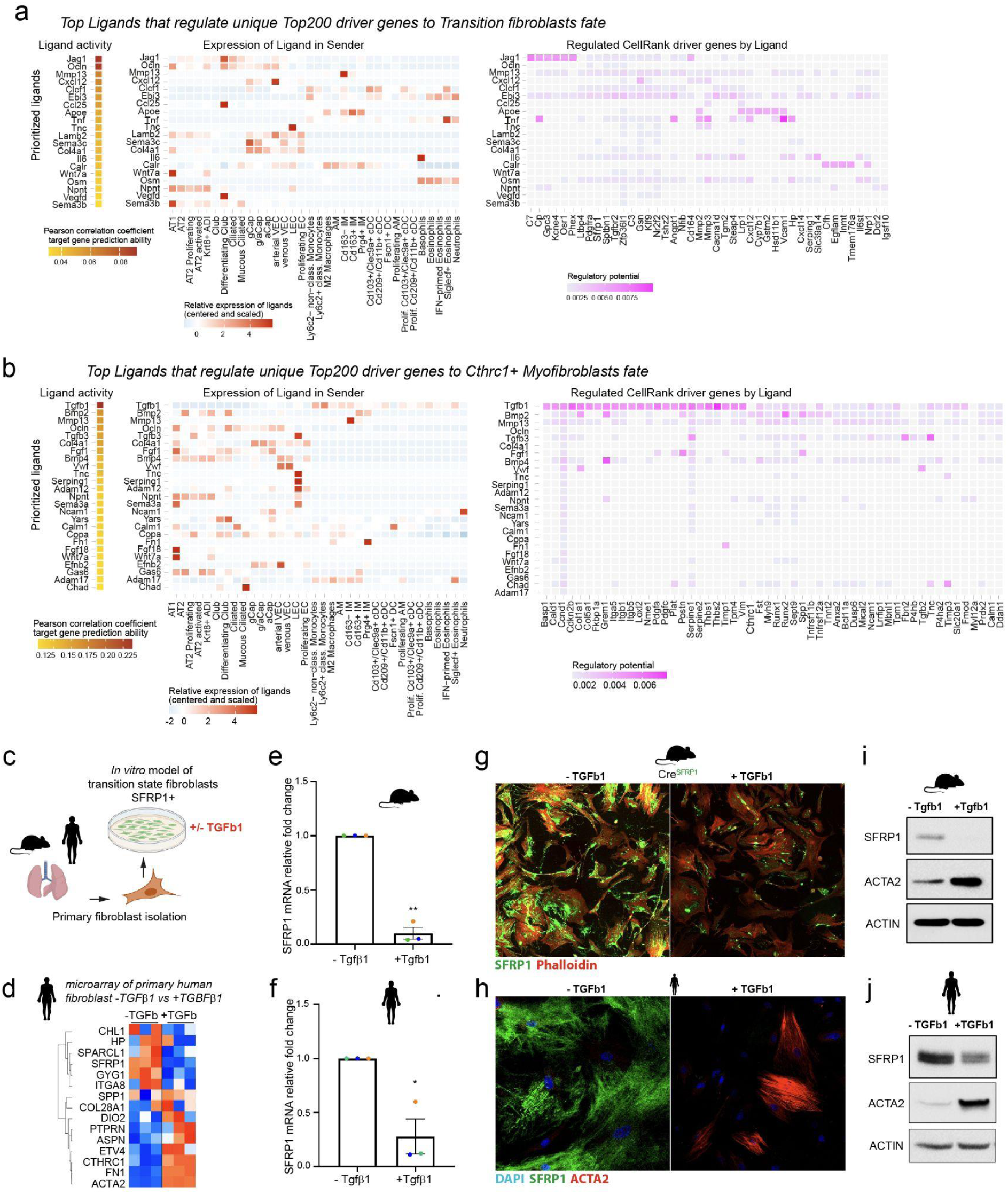
Tgfβ1 drives conversion of Sfrp1+ transitional cells to Cthrc1+ myofibroblasts. **a,b)** NicheNet analysis based prediction of regulatory ligands upstream of fibroblast states. Heatmaps show top ranked ligands with highest potential of regulating the top 200 driver genes towards the fate of **a)** Sfrp1+ transitional fibroblasts and **b)** Cthrc1+ myofibroblasts. The left panel shows scaled expression of those ligands across cell types of the whole lung, while the right panel shows the top downstream target genes of each ligand. **c)** Experimental design. **d)** Heatmap shows z-scored gene expression from bulk transcriptomics after Tgfβ1 treatment of primary human fibroblasts grown in vitro. **e, f)** The bar graph shows expression of Sfrp1 mRNA (q-PCR) 48 hours after TGFβ1-treatment in primary mouse (e) or human (f) lung fibroblasts grown in vitro. Data are expressed as the mean +/− SEM. **P < 0.01, paired two-tailed t-test, n = 3 (fibroblasts from 3 different mice/donors). **g)** Immunofluorescence staining of primary mouse fibroblasts isolated from lungs from a Sfrp1^tm1(EGFP/cre/ERT2)^ mouse. Cultured fibroblasts were treated with Tamoxifen to induce EGFP-expression by the Sfrp1-promoter. 48 hours of Tgfβ1-treatment substantially reduced Sfrp1 levels (right panel in green). Actin filaments were stained with phalloidin (red). Scale bar = 100 µm. **h)** Immunofluorescence staining of primary human lung fibroblasts shows expression of Sfrp1 and Acta2 with and without 48 hours of Tgfβ1-treatment. Scale bars = 50 µm. **i, j)** Expression of the indicated proteins was analyzed using western blotting. Primary mouse (i) and human (j) fibroblasts were cultured in vitro for 48 hours with and without Tgfβ1 stimulation. Representative of n=3.

The highest ranking ligand upstream of the Sfrp1+ transitional cell state was the Notch ligand Jag1, which was highly expressed in secretory airway epithelial cells but also to a lower extent in alveolar epithelial and vascular endothelial cells (Fig. 6a). Indeed, Notch deficiency in mesenchymal cells has been shown to reduce fibrotic remodeling and myofibroblast differentiation in the bleomycin model^32^, which is consistent with our result here. Other top ranked ligands included the Cardiotrophin-like cytokine factor 1 (Clcf1) and the Interleukin-27 subunit beta (Ebi3), which were primarily derived from immune cells of the myeloid lineage, and the granulocyte derived inflammatory factor TNF-alpha (Tnf) (Fig. 6a).

Notably, the highly ranked ligands upstream of Sfrp1+ transitional cell state driver genes did not include Tgfβ, which was however the top ranked ligand for Cthrc1+ myofibroblast driver genes. We found Tgfβ1 as the top ligand, which was mainly expressed by immune cells of the myeloid lineage, as well as Tgfβ3 expressed by activated AT2 cells and lymphatic endothelial cells (Fig. 6b). Other unique ligands upstream of Cthrc1+ myofibroblast driver genes included Bmp2 and Bmp4, which were mostly derived from alveolar epithelial cells, as well as interstitial macrophages in the case of Bmp2 and vascular endothelial cells in the case of Bmp4 (Fig. 6b).

Isolating both mouse and human lung fibroblasts and by culturing them in vitro in the presence of serum, we noted that they exhibited a marker gene expression consistent with the Sfrp1+ transitional state seen in vivo (Fig. 6c, d). Transcriptomes of primary fibroblasts cultured with and without 1 ng/ml Tgfβ1 for 48 hours were characterized by expression of the transition state markers Col28a1 and Sfrp1 in unstimulated conditions, while these genes were downregulated in the Tgfβ1 stimulated condition. Genes denoting myofibroblast identity like Cthrc1 and Spp1, as well as fibrosis-relevant ECM markers like Tnc and Fn1, were upregulated upon Tgfβ1 stimulation (Fig. 6d). We validated these observations in both mouse and human primary lung fibroblasts using real time PCR (Fig. 6e, f), immunofluorescence (Fig. 6g, h), and western blotting (Fig. 6i, j).

In conclusion, our data suggests that the Wnt-modulator Sfrp1 denotes diverse differentiation states of fibroblasts involved in the course of tissue fibrosis, progressing from normal steady state tissue-fibroblasts (Sfrp1^low^), towards an activated transition-state (Sfrp1^high^), and a terminal Cthrc1+/Spp1+ myofibroblast state (Sfrp1^low^) *in vivo* as well as *in vitro*. Our in vitro experiments validate the computational predictions and show that Tgfβ1 functions as a master switch for downregulation of Sfrp1 and upregulation of myofibroblast genes in both mouse and human primary fibroblasts.

### Autocrine signaling by Sfrp1 controls invasion and RhoA activity in Sfrp1+ transitional fibroblasts

Secreted Sfrp1 is described as an inhibitor of the Wnt signaling pathway^33,34^. Additionally, downregulation of Sfrp1 in various cancers was shown to promote tumor cell invasion which directly correlated with poor disease prognosis^35,36^. In the current model of fibrogenesis, activated fibroblasts are thought to migrate into damaged tissue regions, where they transform into myofibroblasts forming so-called fibrotic foci. Surprisingly, transplantation experiments in mouse lungs after bleomycin injury demonstrated that Cthrc1+ myofibroblasts have the highest invasive migratory capacity compared to other fibroblast states^3^.

We have previously identified a transcriptomic invasion signature of collagen-invading lung fibroblasts, which was characterized by a robust reduction in Sfrp1 expression^37^. To directly analyze the association of Sfrp1 expression with the invasive capacity of fibroblasts, we made use of an Sfrp1^tm1(EGFP/cre/ERT2)^ mouse, which after tamoxifen induction expressed EGFP as a fluorescent reporter (Fig. 7a). Using flow cytometry we live sorted Sfrp1^+^-fibroblasts (EGFP-positive) and Sfrp1^−^-fibroblasts (EGFP-negative), and tested their capacity to invade a collagen-I matrix (Fig. 7a, b). Sfrp1^−^-fibroblasts showed a substantially higher invasion capacity than Sfrp1^+^-fibroblasts, respectively (Fig. 7b).

**Figure 7.**
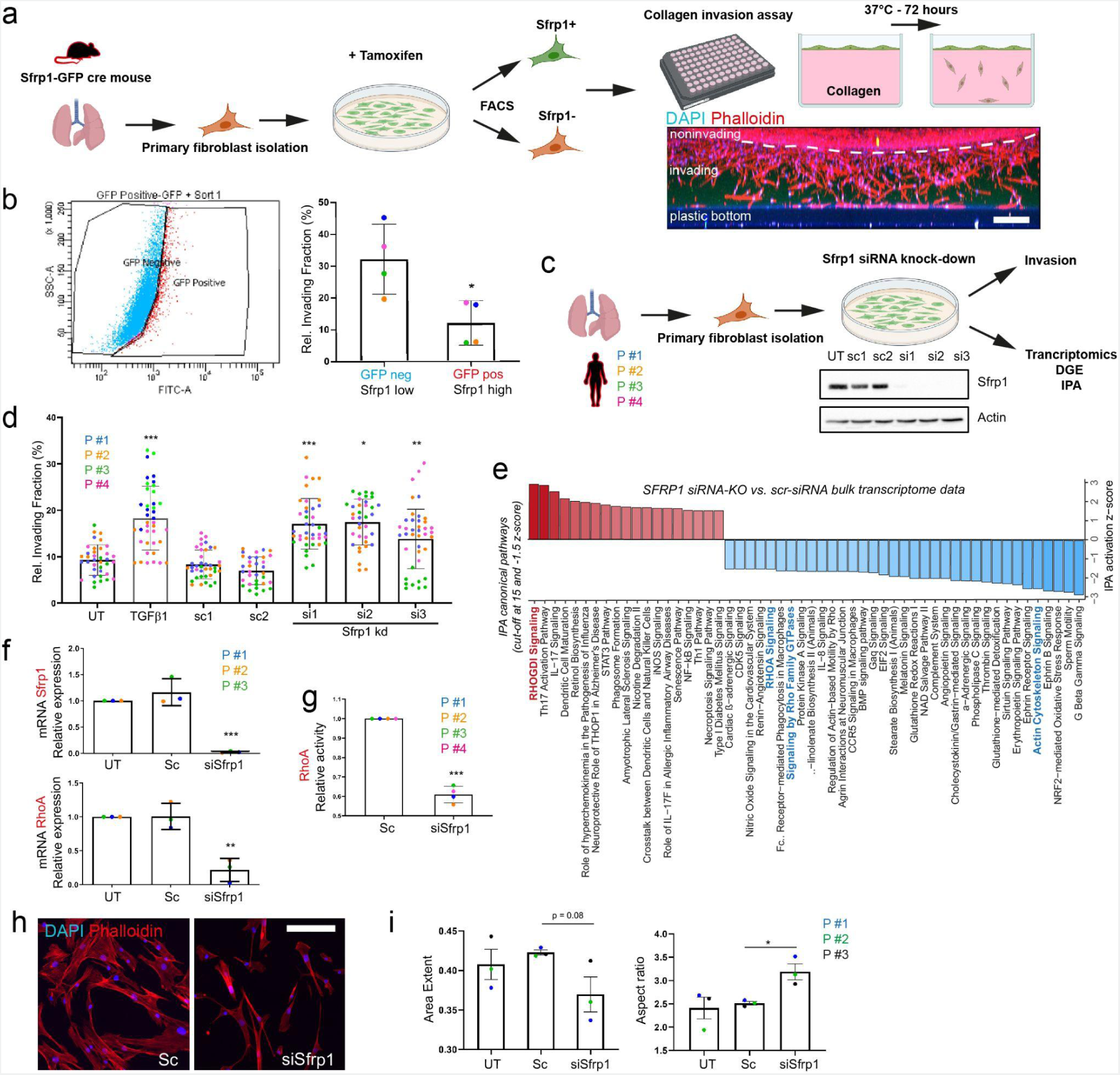
Sfrp1 in transition state fibroblasts regulates invasion as well as RhoA activity and cell shapes. **a)** Primary fibroblasts were isolated from lungs of an Sfrp1^tm1(EGFP/cre/ERT2)^ mouse, cultured on rigid 2D surfaces and subsequently treated with tamoxifen to trigger expression of the fluorescent EGFP reporter indicating Sfrp1 expression. Sfrp1^+^-fibroblasts (EGFP-positive) were separated from Sfrp1^−^-fibroblasts (EGFP-negative) by FACS, and then both subpopulations were applied to an ECM-invasion assay using collagenI as the invasion matrix. Scale bar = 200 µm. **b)** FACS-plot exhibiting separation of Sfrp1^+^-from Sfrp1^−^-fibroblasts. ECM invasion capacity was significantly increased in EGFP-negative Sfrp1^−^-fibroblasts. Data are expressed as the mean +/− SEM. *P < 0.05, paired two-tailed t-test, n = 4 (fibroblasts from 4 different mice). **c)** Primary human fibroblasts were isolated from lungs of human patients and cultured on rigid 2D surfaces. Sfrp1 expression was diminished in these cells by application of three different siRNAs targeting various exons. The successful knock-down of Sfrp1 protein was confirmed by western blot analysis. **d)** Tgfβ1-treatment as well as Sfrp1-depletion augmented the ECM invasion capacity in primary human lung fibroblasts. Data are expressed as the mean +/− SEM. *P < 0.05, **P < 0.01, ***P < 0.001, one-way ANOVA with Dunnett’s multiple comparison test, n = 4 (fibroblasts from 4 different patients). **e)** IPA canonical pathway analysis based on bulk transcriptome data analysis of Sfrp1-siRNA-depleted compared to Sfrp1-expressing primary human lung fibroblasts. IPA identified RhoA-signaling as a highly affected pathway in Sfrp1-siRNA-depleted fibroblasts. **f)** Transcript analysis by qPCR confirmed the successful reduction of Sfrp1 mRNA in Sfrp1-siRNA-treated primary human lung fibroblasts. RhoA mRNA was found to be considerably reduced in Sfrp1-depleted fibroblasts. Data are expressed as the mean +/− SEM. Paired two-tailed t-test. ***P < 0.001, **P < 0.01, n = 3 (fibroblasts from 3 different patients). **g)** Consistent with IPA analysis shown in (e) as well as reduction of RhoA mRNA displayed in (f), RhoA-GTPase activity was significantly reduced in Sfrp1-depleted primary human lung fibroblasts. Data are expressed as the mean +/− SEM. Paired two-tailed t-test. ***P < 0.001, n = 4 (fibroblasts from 4 different patients). **h)** Immunofluorescence labeling of filamentous actin cytoskeleton (Phalloidin in red) indicated a morphological switch towards smaller and elongated cell shapes in Sfrp1-depleted primary human lung fibroblasts. Scale bar = 100 µm. **i)** Detailed cell morphological analysis using an automatized workflow in CellProfiler software. The unbiased quantification of 1000 cells from untreated (UT), scrambled-siRNA-treated (Sc) and Sfrp1-siRNA-treated (siSfrp1) human lung fibroblasts confirmed a substantial switch towards more elongated (smaller area extent, higher aspect ratio) cell morphologies in Sfrp1-depleted primary human lung fibroblasts. n = 1000 fibroblasts per patient. Data are expressed as the mean +/− SEM. Unpaired t-test. P = 0.08, *P < 0.05, n = 3 (fibroblasts from 3 different patients).

To determine if Sfrp1 is mechanistically involved in controlling fibroblast invasion we turned to a loss of function approach using siRNA. We isolated primary human lung fibroblasts from 4 donors and depleted Sfrp1 expression before subjecting these cells to the collagen invasion assay (Fig. 7c). Indeed, primary human lung fibroblasts devoid of Sfrp1 expression considerably increased their invasive capacity compared to untreated and siRNA-scrambled controls (p < 0.001) (Fig. 7d). Furthermore, Tgfβ1-treatment, which we demonstrated to diminish Sfrp1 expression (Fig. 6e-j), caused an increase in the fibroblasts’ invasion capacity compared to untreated controls (p < 0.001)(Fig. 7d).

Bulk transcriptomic analysis of the siRNA-treated fibroblasts revealed significant regulation of the RhoA-signaling-related pathways in Sfrp1-depleted fibroblasts (Fig. 7e). Specifically, using upstream regulator and pathway analysis in the Ingenuity Pathway analysis toolbox we found that “RHOGDI signaling”, an inhibitory pathway for RhoA-activity, was predicted to be highly activated (positive IPA activation z-score), whereas the pathways “RHOA signaling”, “signaling by Rho family GTPases” and “actin cytoskeleton signaling” were found amongst the most deactivated signaling pathways (negative IPA activation z-score) in Sfrp1 knock down cells (Fig. 7e).

Validation by qPCR analysis confirmed the robust reduction of Sfrp1 mRNA in Sfrp1-siRNA-depleted fibroblasts which was concomitant with a reduction of RhoA-mRNA expression by 4.8 fold (P < 0.01) (Fig. 7f). In fact, experimental assessment of RhoA-GTPase activity supported our previous IPA prediction, as RhoA-GTPase activity was found to be diminished about 40% (p < 0.001) in Sfrp1-deficient primary human lung fibroblasts (Fig. 7g). Finally, immunofluorescence labeling of the fibroblasts’ filamentous actin cytoskeleton indicated more elongated cell morphologies in Sfrp1-depleted fibroblasts (Fig. 7h). We used the CellProfiler software for a comprehensive morphological analysis based on actin cytoskeleton stainings and found that Sfrp1 depletion resulted in a 25% decrease of the cell shape parameter “area-extent” and a 30% increase (p < 0.05) in the cells’ aspect ratio, both denoting a switch towards elongated cell shapes (n=1000) (Fig. 7i).

In summary, our loss of function experiments in primary human lung fibroblasts reveal a function of Sfrp1 in regulating fibroblast invasion via the RhoA pathway.

## Discussion

Fibroblasts are master orchestrators of tissue homeostasis, immune reactions, and wound healing throughout the body. Next to the collagen producing smooth muscle cells and pericytes, we found four distinct fibroblast cell types and three distinct injury induced fibroblast activation states that were associated with specific niches in peribronchiolar, adventitial and alveolar locations of the lung. The time-resolved single cell and lineage tracing experiments uncovered highly cell type and lineage specific paracrine signals that likely mediate specialized functions of these fibroblast populations in lung regeneration that await further investigations. Our analysis charts the dynamic evolution of mesenchymal cell states during lung injury and regeneration and reveals autocrine Sfrp1 as a key regulator of fibroblast invasion during the myofibroblastic transition (Fig. 8).

**Figure 8.**
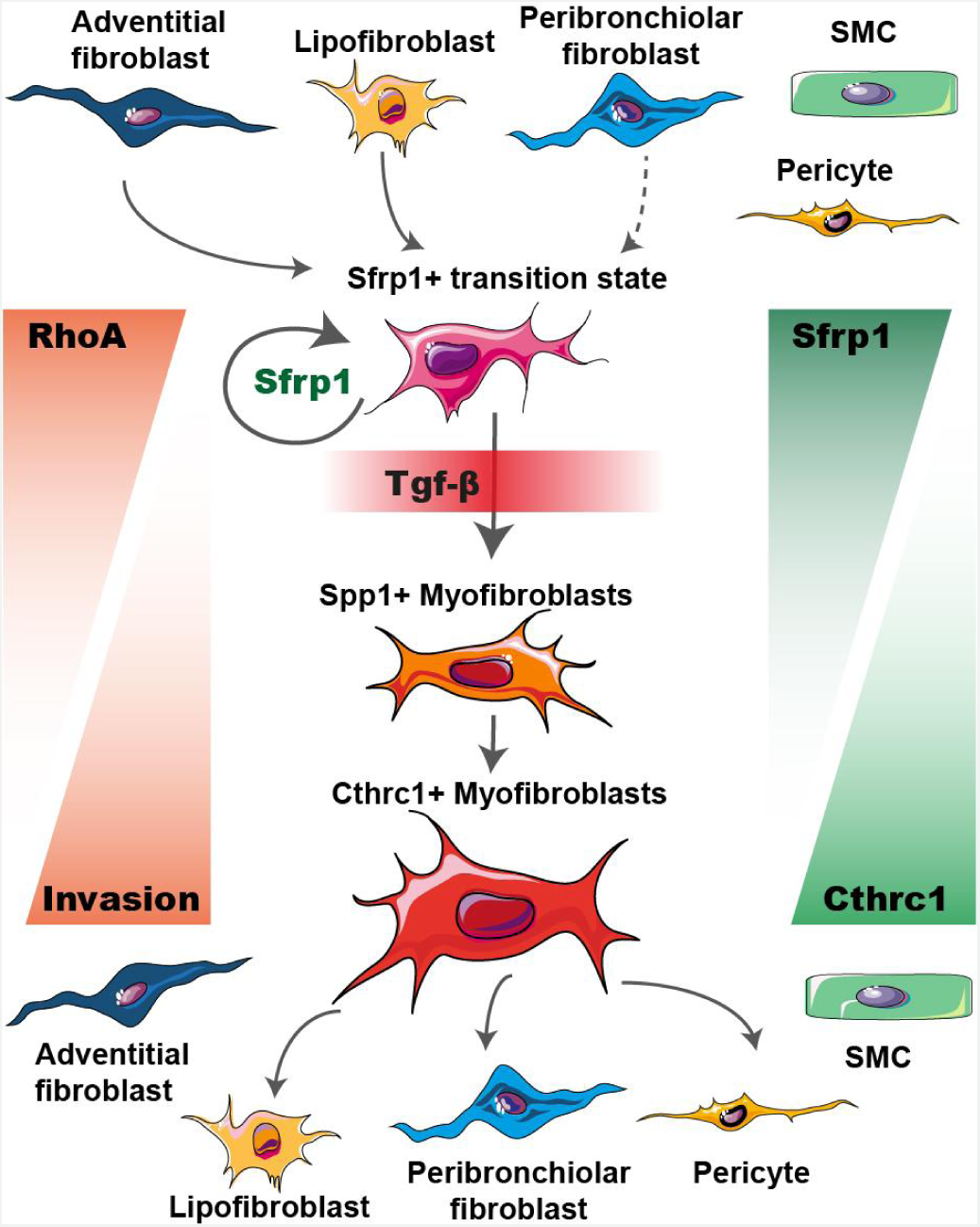
Model illustrating fibroblast phenotypic trajectories in lung repair and regeneration. Our findings reveal the spatiotemporal evolution of distinct fibroblast states after lung injury. We discovered a novel transitional fibroblast state which is characterized by the expression of Sfrp1, as well as its emergence from adventitial, peribronchial and lipofibroblasts early after injury. Autocrine regulation by high levels of Sfrp1 prevents transitional fibroblasts from escaping out of their local niche by invasion. However, in the presence of immune cell-derived Tgfβ1, transitional fibroblasts will start to invade and express the myofibroblast program, indicated by loss of Sfrp1 expression along with the upregulation of myofibroblast marker genes such as Acta2, Spp1 and Cthrc1. Mechanistically, Sfrp1 mediated autocrine signaling inhibits cell invasion and RhoA pathway activity.

The trajectory inference in this study benefits from the very high time resolution after injury (daily sampling) and suggests that adventitial fibroblasts, peribronchiolar and alveolar fibroblasts transcriptionally converge after injury. To our knowledge, this has not been demonstrated before and will require the development of specific genetic lineage tracing tools for experimentally validating our computational predictions. We used single cell transcriptomics on lungs from the Tcf21−Cre reporter mice to address the lineage trajectory of lipofibroblasts. A limitation of these experiments was however that all other mesenchymal cell types with exception of the Hhip+ peribronchiolar fibroblasts were also lineage labeled due to low level expression of Tcf21 in these cells. Tcf21 lineage negative Hhip+ cells were interestingly heavily expanded after injury. We show that these cells can reside in direct physical contact with AT2 cells and are the exclusive source of morphogens such as Wnt5a after injury, which likely has important implications for AT2 progenitor cell function^29^.

Previous work has shown that tissue invasion by fibroblasts is an essential driver of disease progression and severity in lung fibrosis^3,38–40^. We show that the early Sfrp1+ transitional state of injury activated fibroblasts is non-invasive, suggesting that at this stage the activated fibroblasts serve local functions in their niche, such as recruiting immune cells in case of adventitial fibroblasts, or activating epithelial stem cells in case of the lipofibroblasts, pericytes and Hhip+ fibroblasts. Our data indicates that recruited myeloid cells, which heavily accumulate in the first week after injury in the bleomycin model^7^, as well as activated epithelial and lymphatic endothelial cells, produce Tgfβ to switch the non-invasive Sfrp1+ transitional cells into highly invasive Cthrc1+/Spp1+ myofibroblasts. Indeed, a profibrotic cell circuit between macrophages and fibroblasts has been described, which required cadherin-11 mediated direct cell-cell interactions to promote latent Tgfβ activation^41^.

To the best of our knowledge it is currently unclear what the physiological pro-regenerative function of myofibroblast invasion and motility might be. An interesting conundrum is that the classical view of a sessile actomyosin based contractile phenotype of myofibroblasts does not fit our observations that Cthrc1+/Acta2+ myofibroblasts are actually highly invasive and motile. In pathology, such as IPF, the myofibroblasts invade to organize themselves into dense accumulations called fibroblast foci, but also in the bleomycin model they accumulate in clusters via active tissue invasion^39^. Thus, the mechanisms discovered in this study have direct implications for IPF as they may open the way for new therapeutic interventions to limit or reverse fibroblast foci formation.

## Supporting information

Marker gene per cell type for 10x Tcf21 PBS subset

Marker gene per cell type for 10x Tcf21 bleomycin day14 subset

Marker gene per cell type for Dropseq bleomycin timecourse

Spline table for DGE genes across timepoints per cell type

Sequences for SCRINSHOT probes

List of antibodies

Deparaffinization protocol

Sequence of primers used for qRT-PCR studies

## Acknowledgments

We acknowledge the provision of human biomaterial and clinical data from the CPC-M bioArchive and its partners at the Asklepios Biobank Gauting, the Klinikum der Universität München, and the Ludwig-Maximilians-Universität München. We are grateful to M. Neumann and A. van den Berg from the Comprehensive Pneumology Center (CPC) for providing superb technical support. We also thank Dr. Inti Alberto de la Rosa Velazquez and team from the genomics core facility of Helmholtz Munich for expert sequencing service.

We acknowledge support by the German Center for Lung Research (DZL), the Helmholtz Association, the European Union’s Horizon 2020 research and innovation program (grant agreement 874656) and the Chan Zuckerberg Initiative (CZF2019-002438).

## Author contributions

Conceptualization and supervision: G.B. and H.B.S.; Methodology and investigation: C.H.M., A.S., M.A., J.C.P., P.O., I.A., A.L., A.R.-C., N.J.L., M.S., S.A., D.P.-G., B.O., V.V.-A., M.I., M.G.; Data analysis: I.E.-F., M.T., M.I., J.B., O.E., A.O.Y., K.A., R.E.M., C.S., F.J.T., G.B., H.B.S; Surgical work and human tissue: G.M.S, J.Behr, N.K.; Writing: C.H.M., A.S., G.B., H.B.S.

## Supplementary figures

**Figure S1.**
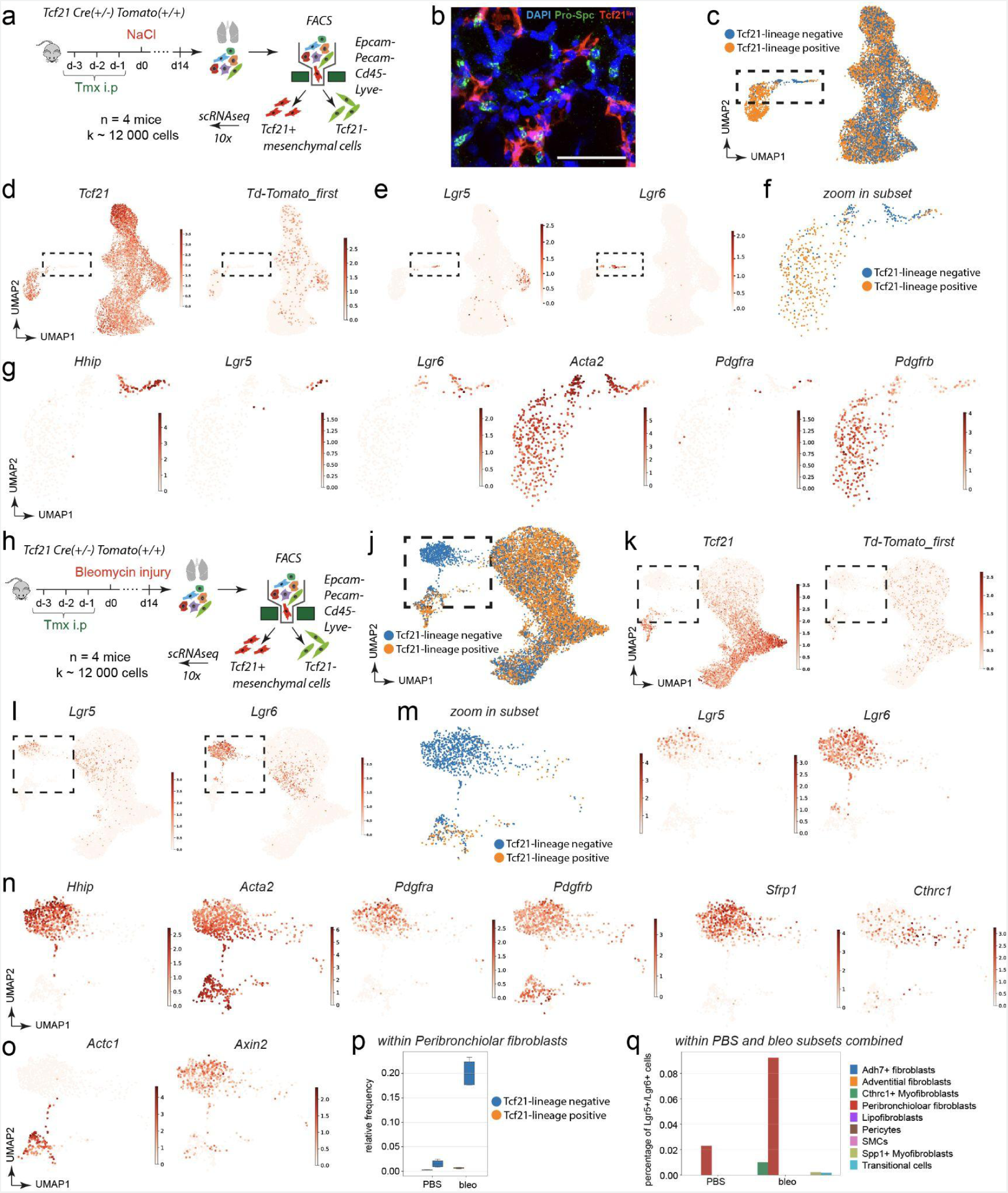
Complexity within Hhip+ peribronchiolar fibroblast population. **a,h)** Adult Tcf21−Cre(+/−)-Td-Tomato(+/−) were injected with Tamoxifen three days before applying **a)** NaCl or **h)** bleomycin and sacrificed at day 14. Isolated cells from mouse lungs were negative selected (Epca−/Pecam−/Cd45/Lyve−) for mesenchymal cells using FACS, and Tcf21 lineage positive and negative cells were sorted and subjected for scRNAseq analysis using the 10x Genomics platform. **b)** Immunofluorescent analysis of Pro-SPC (green) shows AT-2 cells and Tcf21+Td-tomato+ (red) positive mesenchymal cells in the lung. **c, j)** UMAP embeddings display cells colored by Tcf21 lineage identity for c) PBS or j) bleomycin derived cells respectively, **d,k)** Tcf21 and Td-tomato gene expression, **e,i)** Lgr5 and Lgr6 expression. Rectangle indicates subset of cells used for the embedding panels in f-g. **f-g)** Zoom into PBS derived subset of Lgr5+/Lgr6+ positive cells shows UMAP embendings displaying Hhip, Lgr5, Lgr6, Acta2, Pdgfra and Pdgfrb gene expression (g). **m-o)** Zoom into bleomycin day14 derived subset of Lgr5+/Lgr6 positive cells shows UMAP embendings displaying **m)** Lgr5 and Lgr6, **n)** Hhip, Acta2, Pdgfra, Pdgfrb, Sfrp1, Cthrc1, and **o)** Actc1 and Axin2 gene expression. **p)** Frequency analysis of Tcf21 lineage positive and negative cells within the population of Peribronchiolar fibroblasts. **q)** Frequency analysis of Lgr5+/Lgr6+ positive cells across all mesenchymal subtypes and between PBS and bleomycin condition.

**Figure S2.**
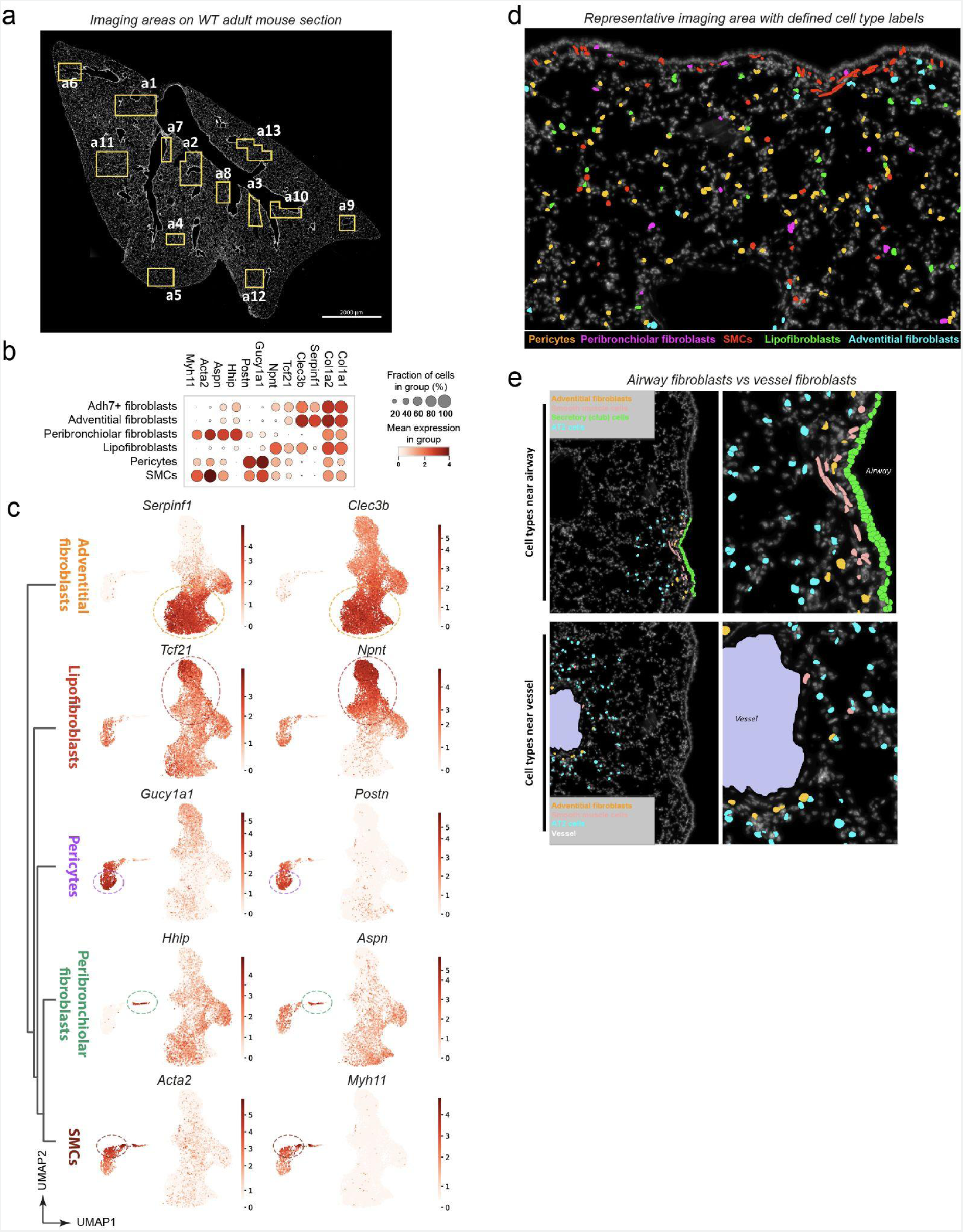
SCRINSHOT analysis of mesenchymal subtypes. **a)** Regions shown were used for SCRINSHOT on a healthy mouse lung cryo-section. **b)** Dotplot shows gene expression of the two metacelltype marker genes that were chosen for SCRINSHOT analysis. **c)** UMAP plots depict gene expression of the two metacelltype marker genes that were chosen for SCRINSHOT analysis for each mesenchymal cell type. **d)** Image depicts representative region of the mouse lung with cells labeled based on their SCRINSHOT marker expression. **e)** Images show representative regions for airway and vessel domains. **f)** Raw images show interaction of mesenchymal cell types with AT-2 cells in alveolar regions.

**Figure S3.**
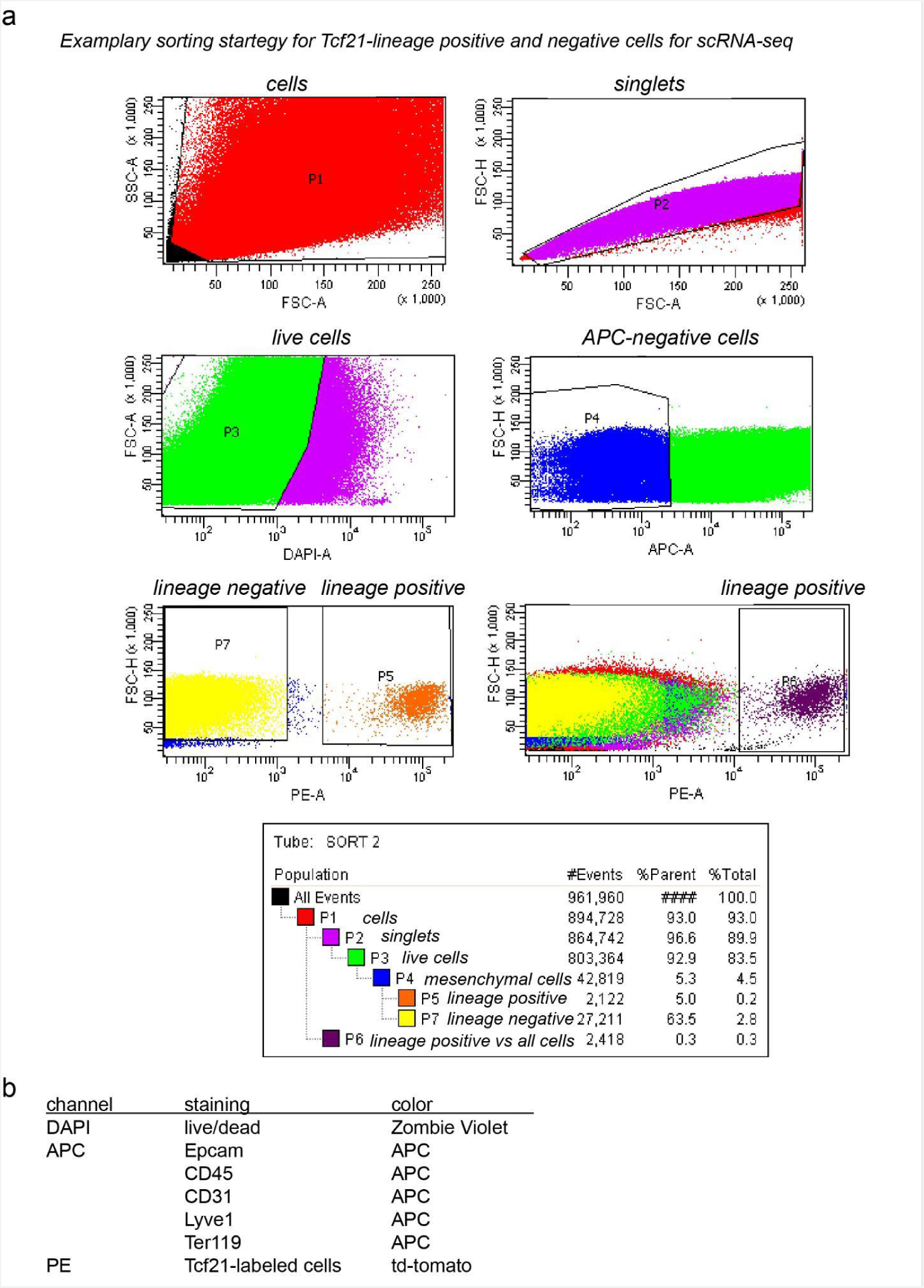
Gating strategy for Tcf21 lineage positive and negative mesenchymal cells. **a)** Exemplary gating strategy for the analysis of Epcam−/CD45−/CD31−/Lyve−/Ter119− negative selected, living mesenchymal cells from Tcf21−Td-tomato mice, that were split into lineage positive (gate P5) and lineage negative cells (gate P7). **b)** List of channels, stained markers and colors used for the sorting.

**Figure S4.**
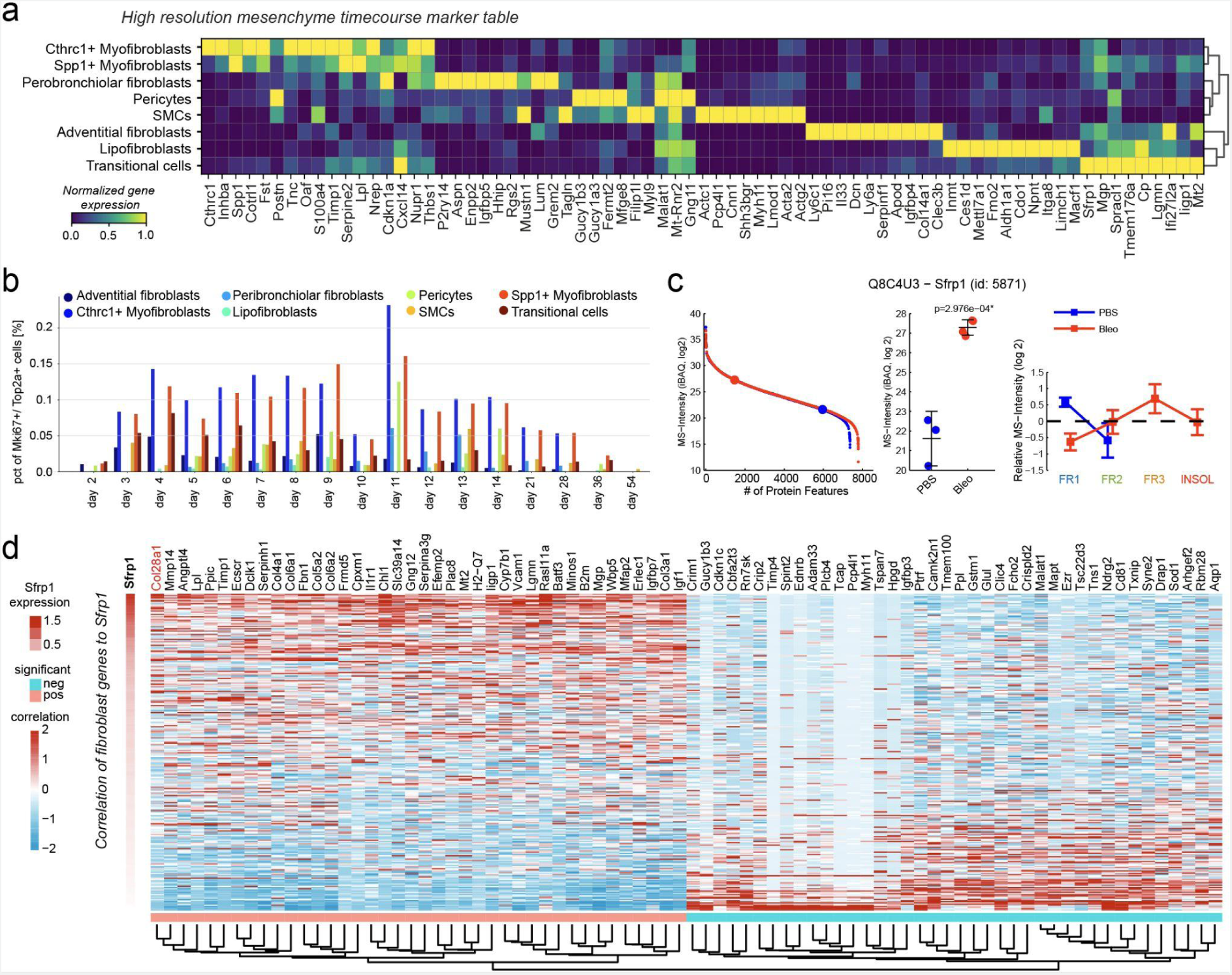
Bleomycin time course analysis. **a)** Matrixplot of marker genes for the high temporal resolution bleomycin time course of mouse lung mesenchymal cell types. **b)** Proliferation analysis shows the percentage of cells expressing Mki67 or Top2a per cell type and time point. **c)** Quantitative proteome data depict the protein abundance rank of Sfrp1 relative to all other quantified proteins, the iBAQ-normalized mean MS-intensity between healthy PBS treated control mice and Bleomycin treated mice from day 14, and the QDSP profiles indicative for the detergent solubility. The mean and standard error of the mean are shown (day 3, *n* = 3; day 14, *n* = 7; day 28, *n* = 4; day 56, *n* = 3) ^25^. **d)** The heatmap shows the expression of genes most strongly associated with Sfrp1 expression within the mesenchymal cells (bottom colorbar indicates significant correlation or anti-correlation).

**Figure S5.**
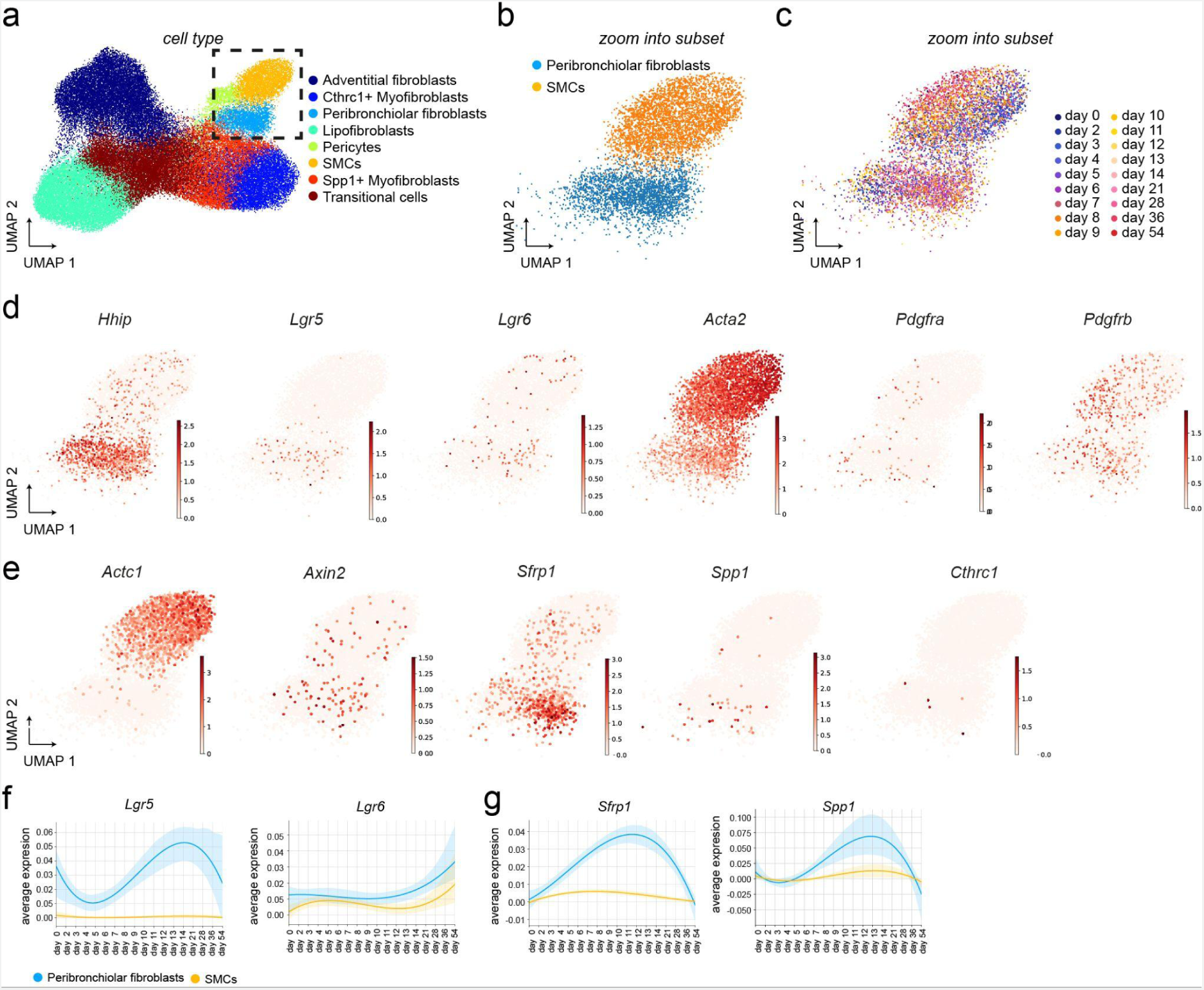
Sub-clustering/zoom in on Hhip+ Peribronchiolar fibroblasts from time course data. **a)** UMAP embeddings display cells colored by cell type identity. Rectangle indicates zoom into a subset of Peribronchiolar fibroblasts and Smooth Muscle Cells (SMCs). **b-c)** UMAPS embeddings display subset of cells colored by **b)** cell type identity **c)** time point and gene expression of **d)** Hhip, Lgr5, Lgr6, Acta, Pdgfra, Pdfgrb as well as **e)** Actc1, Axin2, Sfrp1, Spp1 and Cthrc1. **f-g) T**he line plots show the smoothed mean expression over time for the indicated genes within the subset of Peribronchiolar fibroblasts And SMCs with a confidence interval across the mouse replicates.

**Figure S6.**
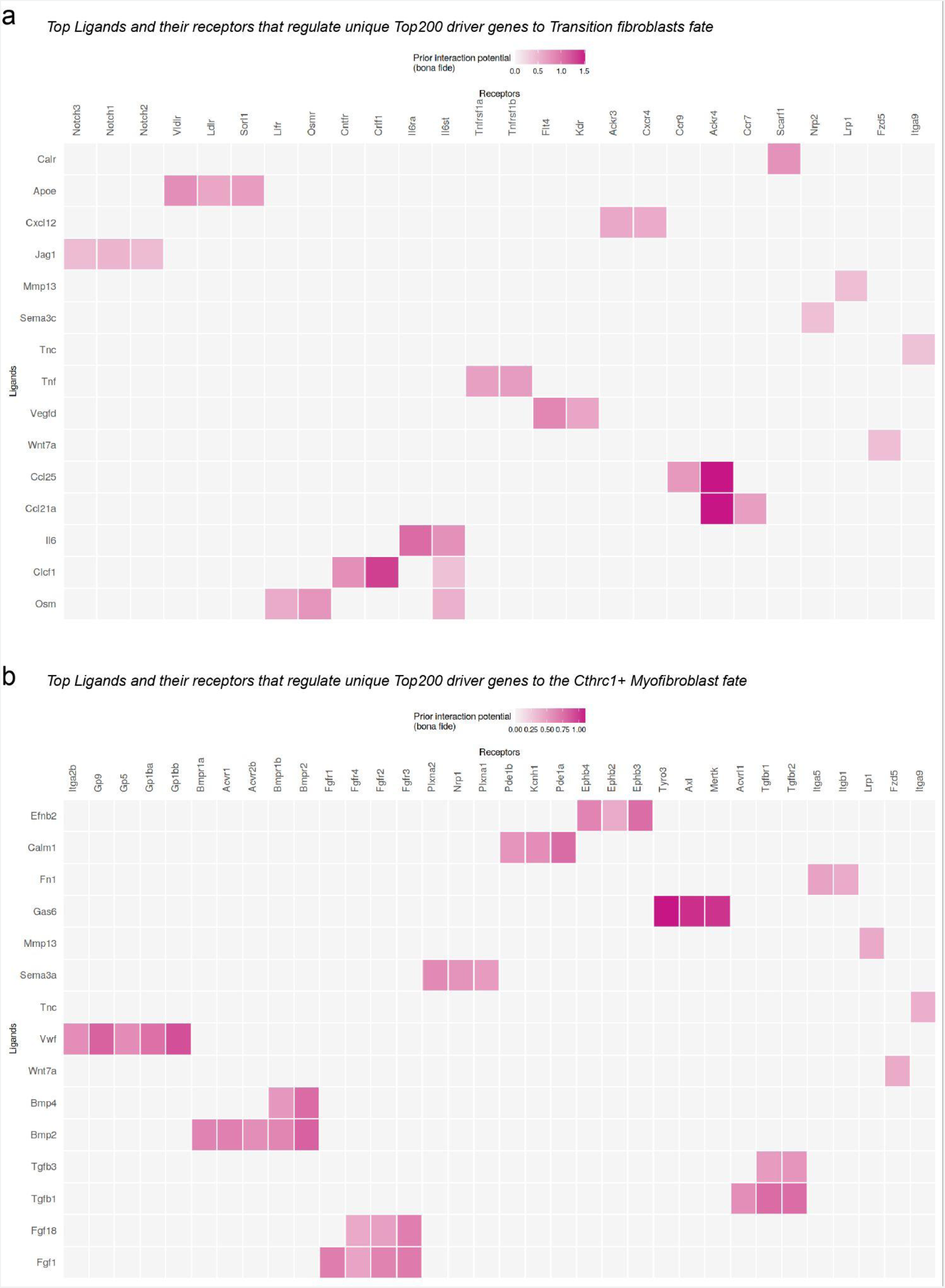
Ligand-receptor pairs. **a-b)** Heatmaps show the regulatory potential for the given ligand and the potential bona fide receptors in a) the Transition fibroblast fate and b) the Cthrc 1 Myofibroblast fate.

**Figure S7.**
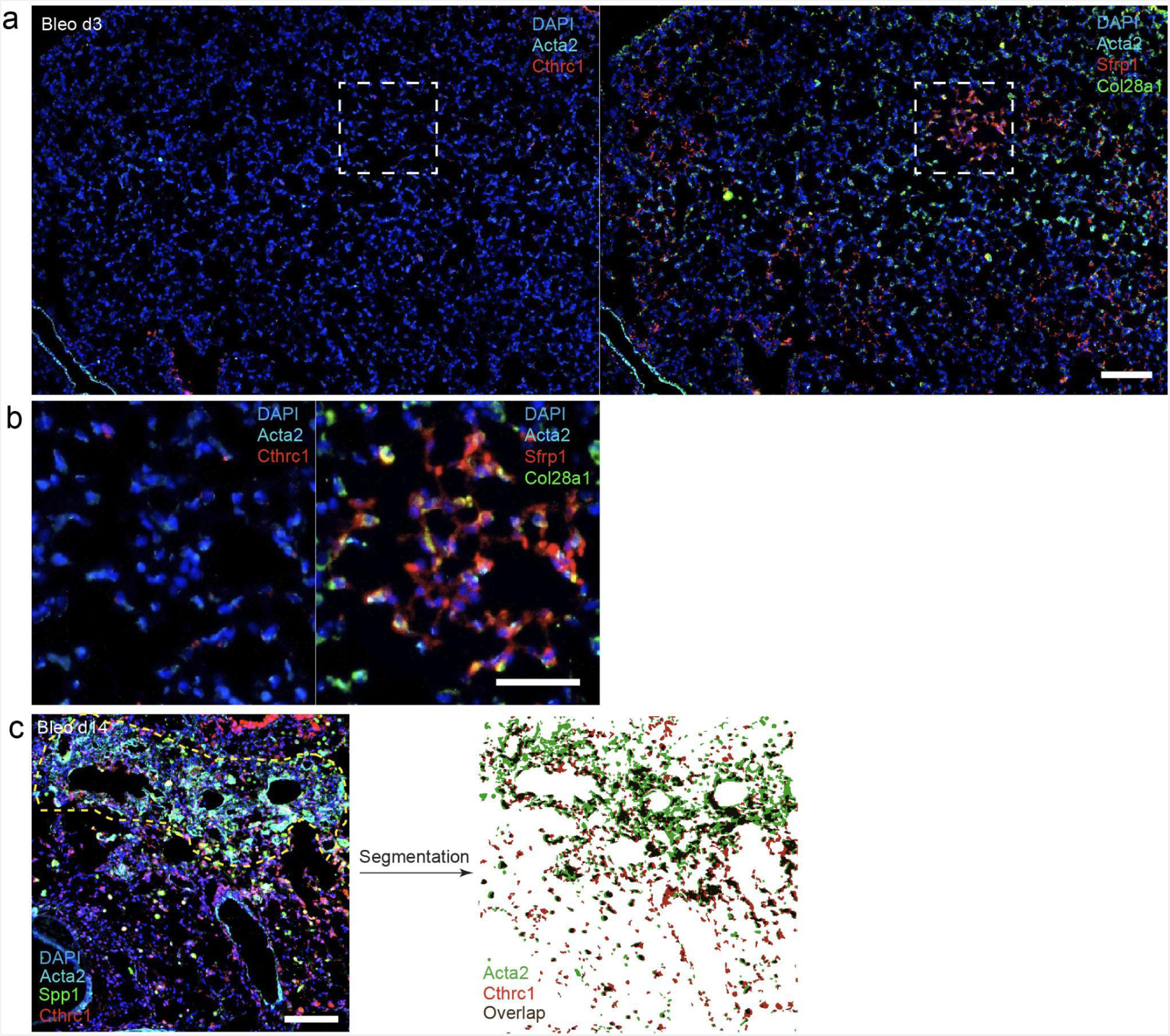
Sfrp1+ transitional fibroblasts emerging in parenchymal lung regions and image segmentation. **(a)** Iterative staining (4i) of parenchymal lung tissue at day 3 after injury indicating emergence of Col28a1 (green)/Sfrp1 (red) double positive cells in the left image. The same cells and regions remained negative for Cthrc1 (red) and Acta2 (cyan). Scale bar = 100 µm. Dashed box indicates magnified view, shown in **(b).** Iterative staining (4i) of parenchymal lung tissue at day 3 after injury indicating Col28a1 (green)/Sfrp1 (red) double positive cells in the left image. The same cells, as indicated by white dashed lines, were found to be Cthrc1 (red) and Acta2 (cyan) negative. Scale bar = 50 µm. **(c)** Representative image demonstrating segmentation which was used for the quantification of appearance of Acta2, Cthrc1, Spp1, Col28a1 and Sfrp1 positive cells at various time points after bleomycin treatment. Yellow dashed-line indicates a densely fibrotic region. Cell nuclei stained with DAPI in blue, Acta2 in cyan, Spp1 in green and Cthrc1 in red. The left panel shows software-based segmentation of Acta2 (green) and Cthrc1 (red). Overlapping segmented objects are depicted in brown. Scale bar = 100 µm.

## Methods

### Data availability

ScRNA-seq data was deposited to the Gene Expression Omnibus database. The high temporal mesenchymal enriched Dropseq can be found with the accession code GSE207851, and the Tcf21 labeled mesenchymal 10x data with the accession code GSE207687. Microarray data of primary human fibroblasts (SFRP1-siRNA knockdown) can be found with the accession code GSE207561.

### Human tissue and ethics statement

Human tissue has been obtained from the Comprehensive Pneumology Center cohort of the BioArchive CPC-M at the University Hospital Grosshadern of the Ludwig Maximilian University (Munich, Germany) and by the Asklepios Biobank of Lung Diseases (Gauting, Germany). Participants provided written informed consent to participate in this study, in accordance with approval by the local ethics committee of the LMU, Germany (Project 333-10, 454-12).

### Animals

Pathogen-free female C57BL/6J mice were purchased from Charles River Germany and Tcf21iCre(+/−);R26tdTomato(+/−) were maintained in the AG Morty at the Max-Planck-Institut für Herz- und Lungenforschung in Bad Nauheim. Animals were maintained at the appropriate biosafety level at constant temperature and humidity with a 12 hour light cycle. Animals were allowed food and water ad libitum. Animal handling, bleomycin/PBS administration, and organ withdrawal were performed in accordance with the governmental and international guidelines and ethical oversight by the local government for the administrative region of Upper Bavaria (Germany), registered under ROB-55.2-2532.Vet_02-16-208.

### Experimental design and bleomycin treatment

Mice were randomly divided into two groups: saline-only (PBS), or bleomycin (Bleo). Lung injury and pulmonary fibrosis were induced by single-dose administration of bleomycin hydrochloride (Sigma Aldrich, Germany), which was dissolved in sterile PBS and given at 2U/kg bodyweight by oropharyngeal instillation. The control group was treated with sterile PBS only. Mice were randomly sacrificed at designated time points (days 2-14, 21, 28, 35, 56) after instillation. Treated animals were continuously under strict observation with respect to abnormal behavior and signs of body weight loss. Tcf21iCre(+/−);R26tdTomato(+/−)-lineage labeled mice, both female and male, were treated with 100mg/kg Tamoxfien 3 days before the start of the bleomycin treatment (3.5U/kg) or NaCl and sacrificed at day 14.

### Generation of single cell suspensions from whole mouse lung tissue

Lung single cell suspensions were generated as previously described ^42^. Briefly, after euthanasia, the mouse lung was perfused with sterile saline through the heart and all lobes were removed, minced, and transferred for mild enzymatic digestion for 20-30 min at 37°C in an enzymatic mix containing dispase (50 caseinolytic U/ml), collagenase (2 mg/ml), elastase (1 mg/ml), and DNase (30 µg/ml). Cells were collected by straining the digested tissue suspension through a 40 micron mesh and red blood cell lysis (RBC Lysis Buffer, ThermoFisher) was performed for 1 min. After centrifugation at 300 x g for 5 minutes, single cells were taken up in 1 ml of PBS supplemented with 10% fetal calf serum, counted and critically assessed for single cell separation and overall cell viability.

### Negative selection by Magnetic-activated cell sorting (MACS)

For mesenchymal cell enrichment by negative MACS selection, cells were stained with 10 µl per 10 million cells of CD326-AlexaFluor647 antibody (Biolegend, 118212), CD31-APC (Invitrogen, 17-0311-82), CD45−APC (Biolegend, 103112), Lyve1-AF647 (Invitrogen, 50-0443-82) and Ter119-APC (Biolegend, 116218) for 30 min at 4°C in the dark. After washing, the labeled cells were incubated with 100 µl per 10 million cells of MACS dead cell removal beads (Miltenyi Biotec, 130-090-101) and 20 µl of each microbeads specific against AlexaFluor647 (Miltenyi Biotec, 130-091-395) and APC (Miltenyi Biotec, 130-090-855) for 15 min at 4°C. MACS LS columns (Miltenyi Biotec, 130-042-401) were prepared according to the manufacturer’s instructions. Cells were applied to the columns and positively-labeled cells, bound by the microbeads, were retained in the magnetic column, while the flow-through containing the negatively enriched mesenchymal cells was collected on ice and applied to the Dropseq workflow.

### Lineage and negative selection of Tcf21 cells by Fluorescence-activated cell sorting (FACS)

Cells were counted and stained with anti-mouse Epcam CD326-AlexaFluor647 antibody (1:100, Biolegend, 118212), CD31-APC (1:250, Invitrogen, 17-0311-82), CD45−APC (1:250, Biolegend, 103112), Lyve1-AF647 (Invitrogen, 50-0443-82), Ter119-APC (1:100, Biolegend, 116218), and Zombie Violet for viability (BioLegend, 423113). All stainings were performed per 10 MIo cells in 1ml. Cells were stained with Zombie violet according to manufacturer’s instructions for 15 min at RT, washed once with PBS and blocked with anti-mouse FC block for 10min on ice. Staining occurred for 20min at 4°C in the dark in the FACS buffer (PBS + 0.1% BSA), and cells were washed once and filtered with a cell strainer (70µM) before subjecting them to FACS sorting. Mesenchymal cells were sorted as ZombieViolet−/APC− and TCF21 lineage was distinguished by the dt-Tomato signal in the PE channel. Data acquisition was performed in a BD Fortessa flow cytometer (Becton Dickinson, Heidelberg, Germany). 30.000 cells per lineage and mouse were sorted into cooled RPMI media and subject to standard 10x Genomics workflow.

### Single cell RNA-sequencing using the Dropseq method

Dropseq experiments were performed according to the original protocols and as previously described^42^. For Dropseq, cells were aliquoted in PBS supplemented with 0.04% of bovine serum albumin at a final concentration of 100 cells/µl. Using the microfluidic device, single cells (100/µl) were co-encapsulated in droplets with barcoded beads (120/µl, purchased from ChemGenes Corporation, Wilmington, MA) at rates of 4000 µl/hr. Droplet emulsions were collected for 10-20 min/each prior to droplet breakage by perfluorooctanol (Sigma-Aldrich). After breakage, beads were harvested and the hybridized mRNA transcripts reverse transcribed (Maxima RT, Thermo Fisher). Unused primers were removed by the addition of exonuclease I (New England Biolabs), following which, beads were washed, counted, and aliquoted for pre-amplification (2000 beads/reaction, equals ca. 100 cells/reaction) with 12 PCR cycles (Smart PCR primer: AAGCAGTGGTATCAACGCAGAGT (100 μM), 2x KAPA HiFi Hotstart Ready-mix (KAPA Biosystems), cycle conditions: 3 min 95°C, 4 cycles of 20s 98°C, 45s 65°C, 3 min 72°C, followed by 8 cycles of 20s 98°C, 20s 67°C, 3 min 72°C, then 5 min at 72°C)^27^. PCR products of each sample were pooled and purified twice by 0.6x clean-up beads (CleanNA), following the manufacturer’s instructions. Prior to tagmentation, complementary DNA (cDNA) samples were loaded on a DNA High Sensitivity Chip on the 2100 Bioanalyzer (Agilent) to ensure transcript integrity, purity, and amount. For each sample, 1 ng of pre-amplified cDNA from an estimated 1000 cells was tagmented by Nextera XT (Illumina) with a custom P5-primer (Integrated DNA Technologies). Single-cell libraries were sequenced in a 100 bp paired-end run on the Illumina HiSeq4000 using 0.2 nM denatured sample and 5% PhiX spike-in. For priming of read 1, 0.5 μM Read1CustSeqB was used (primer sequence: GCCTGTCCGCGGAAGCAGTGGTATCAACGCAGAGTAC).

### Single cell RNA-sequencing using the 10x Genomics method

The sorted cells were were diluted to 1000 cells/µl, and loaded on the 10× Chromium Next GEM Chip G with a targeted cell recovery of 10,000. The following steps were completed according to the manufacturer’s protocol (Chromium Next GEM Single Cell 3ʹ Reagent Kits v3.1). Libraries have been pooled according to their minimal required read counts (35,000 or 50,000 reads/cell for 3′ gene expression libraries, 20,000 reads/cell for 5′ gene expression libraries, and 5000 reads/cell for TCR libraries). Illumina paired-end sequencing was performed with 150 or 200 (3′ gene expression) on a NovaSeq 6000.

### Processing of the high-temporal resolution bleomycin mesenchymal data set

For the high resolution mesenchymal lung data set, the Dropseq computational pipeline was used (version 2.0) as previously described^42^. Briefly, STAR (version 2.5.2a) was used for mapping^43^. Reads were aligned to the mm10 reference genome (provided by the Dropseq group, GSE63269). Downstream analysis was performed using the scanpy package^56^ (v1.6.0). The top 5000 cells with the highest number of transcripts were read in per sample and subjected to quality control. We assessed the quality of our libraries and excluded barcodes with less than 200 genes detected and retained those with a number of transcripts between 200 to maximal 6000. A high proportion (> 10%) of transcript counts derived from mitochondria-encoded genes may indicate low cell quality, and we removed these unqualified cells from downstream analysis.

To mitigate effects of background mRNA contamination, we employed the R library SoupX^44^. We manually set the contamination fraction to 0.3 and corrected the count matrices with adjustCounts(). The expression matrices were normalized with scran’s size factor based approach^45^ and log transformed via scanpy’s pp.log1p() function. Variable genes were selected sample-wise (flavor = “cell_ranger”), excluding known cell cycle genes. Those genes being ranked among the top 4,000 in at least 6 samples were used as input for principal component analysis (8,209 genes). To filter the data further, the cells were clustered and clusters expressing no mesenchymal cell marker Col1a1 were excluded from the data set. The aligned bam files were used as input for Velocyto^46^ to derive the counts of unspliced and spliced reads in loom format. Next, the sample-wise loom files were combined, normalized and log transformed using scvelos (https://github.com/theislab/scvelo) functions normalize_per_cell() and log1p(). After merging the loom information to the exported .h5ad file using scvelos merge() function the object was scaled and the neighborhood graph constructed with Batch balanced KNN (BBKNN)^47^ to account for the different PCR cycles used in the experiment with neighbors_within_batch set to 15 and n_pcs to 40. Several HVGS genes causing an artificial split up of clusters were removed manually. Two dimensional visualization and clustering was carried out with the scanpy functions tl.louvain() at resolution two and tl.umap().

*Time course differential expression analysis*: to identify genes that show differential expression patterns across time within a given cell type we performed the following analysis (already described^42^). We used the R packages splines and lmtest for our modeling approach. Within each cell type we modeled gene expression as a binomial response where the likelihood of detection of each gene within each mouse sample was the dependent variable. Therefore, the sample size of the model was the number of mouse samples (*n* = 28) and not the number of cells. To assess significance we performed a likelihood-ratio test between the following two models. For the first model, the independent variables contained an offset for the log-transformed average total UMI count and a natural splines fit of the time course variable with two degrees of freedom. The independent variables of the second model just contained the offset for the log-transformed average total UMI count. The dependent variable of both models was the number of cells with UMI count greater than zero out of all cells for a given cell type and mouse sample.

### Trajectory inference with CellRank

To model the cell state dynamics within the mesenchymal cells, the CellRank algorithm^26^ was used. The time course single cell data was separated into three distinct time frames day2-day5, day6-d21, and day28-day54 and each was processed separately. On the subsetted embeddings, the velocity information recovered, and Cellrank initiated by defining the combined kernel with weighing the velocity kernel *0.8 and the connectivity kernel wit *0.2, before computing the sure components and macrostates on the cell type level, until each cell type was represented by at least one macrostate; multiple macrostates per cell type were combined into one. Terminal cell states were set for each time frame separately as shown in the figure and the absorption probabilities were calculated. For each terminal state, the lineage drivers were computed.

### Cell-cell communication analysis with NicheNet

NicheNet analysis was used to identify putative upstream regulator and ligand-receptor interactions regulating trajectories towards the transitional and Cthrc1+ myofibroblast state using the R package nichenetr^31^ (version 1.0). To analyze niche signals that drive trajectories towards these two states, we defined them as receiver cells and all cell types contained in a whole mouse lung gene expression dataset from bleomycin and PBS mice at day 14 generated with 10x Chromium Next GEM Single Cell 3ʹ Reagent Kits v3.1 (data not shown) as sender cells.

To define a geneset of interest we used CellRank to compute the top 200 unique lineage driver genes towards the respective cell states. Next, we identified a list of potentially active ligands based in sender cells (% expressed > 10%) and ranked these potential ligands according to their ability (Pearson’s correlation coefficient) to predict the gene set of interest. We then selected the top 25 ligands for subsequent analysis and visualization.

### Processing of the Tcf21−lineage labeled mesenchymal 10x data

After sequencing, the processing of next-generation sequencing reads of the scRNA-seq data was performed using CellRanger version 3.1.0 (10× Genomics) with a modified genome, by using the mus musculus reference genome (mm10) and manually adding the sequence of the dt-Tomtato gene. We performed unsupervised analysis for multiple subsets of the data with scanpy^56^ (v1.6.0). Quality control was performed similarly as described for the Drop-seq data set. We first filtered genes that were expressed in at least 10 cells, scaled cell-wise expression vectors to a total count of 10,000, and logp1-transformed the data and selected highly variable genes (flavor = ”cell_ranger”). Barcodes with less than 250 genes detected and a mitochondrial fraction of less than 10% were excluded, and those with a number of transcripts between 500 to maximal 50 000 were retained. Again, the expression matrices were normalized with scran’s size factor based approach^45^ and log transformed via scanpy’s pp.log1p() function. Variable genes were selected sample-wise (flavor = “cell_ranger”) and known cell cycle genes were excluded. Those genes being ranked among the top 4,000 in at least 3 samples were used as input for principal component analysis (8,162 genes). We then performed a principal component analysis, based on which we computed the k-nearest neighbor graph (n_pcs = 40, n_neighbors = 20). For two-dimensional visualization we used tl.umap()^57^. Clustering was established using the Leiden and Louvain algorithm^58^.

### Multiplexed in situ hybridization - SCRINSHOT

SCRINSHOT experiments were performed to an earlier version of the method, now published ^23^. Briefly, after euthanasia, mouse lungs were washed with PBS and immediately inflated with 4% paraformaldehyde. For frozen sections, tissue was fixed for 1h at room temperature, embedded in OCT media on dry ice and stored at −80°C. Thin lung sections (7 µm) were cut on a cryostat. Multiplexed in situ hybridization was performed as described previously. Briefly, frozen sections and their cells were permeabilized with 0.1 M HCl in DEPC water at room temperature for 3 minutes and dehydrated in an ethanol series. An incubation chamber was mounted and nonspecific mRNA binding was blocked by adding oligo-dT fragments. Three to five gene specific hybridization probes (padlock probes) were added for 5min at 55°C and for 90 min at 45°C. Ligation of the circular padlock probes was ensured by incubation with SplitR enzyme at room temperature overnight. Rolling circle amplification (RCA) of the padlock circle at 30°C overnight followed by a degradation step for non-specific products with uracil-N-glycosylase (UNG) for 45 minutes at 37°C, provided an amplified number of binding sites for labeled detection probes. One specific labeled probe per padlock probe, with probes for the same gene sharing one color, were hybridized to their binding sites for 60 min at room temperature in the dark. After washing, the tissue was hydrated, the incubation chamber removed and the slide mounted with a cover slip for microscopy. Per hybridization cycle, three genes and DAPI could be visualized. After successful imaging, the cover slip and mounting media were removed, the tissue dehydrated again, and the procedure repeated from the UNG step onwards with different labeled probes. Probes used in the experiment were directed against: Postn, Gucy1a3, Aspn, Hhip, Myh11, Acta2, Tcf21, Serpinf1, Clec3b, Col1a2, Rtkn2, Scgb1a1, Sftpc, Clalcr. Respective sequences for all padlock and detection probes can be found in Table S5.

### Cell culture and treatments

Primary human fibroblasts (phFbs) and primary mouse fibroblasts (pmFbs) were cultivated in 24-well plates with a density of 1.0 ×10^5^ cells/ well and in 6-well plates with a cell density of 2.5×10^5^ cells/ well. DMEM-F12 medium supplemented with 20 % FBS and 100 U/ml of penicillin/streptomycin was used. For Tgfβ1 treatment, cells were initially synchronized using starvation medium (DMEM-F12, 1% FBS and 100 U/ml penicillin/streptomycin) for 24 hours prior to any instillation. Subsequently, cells were induced with 1 ng/ml of human recombinant Tgfβ1 (R&D) for 48 hours. Cells were then harvested for protein and mRNA isolation. phFbs were treated with human siRNAs for Sfrp1. siRNAs obtained from Thermo Fisher Scientific were initially resuspended in nuclease-free water to a final concentration of 2µM to prepare a stock solution. For transfection, 10nM final concentration of Sfrp1 siRNA was added to 5µl of Lipofectamine RNAiMax (Invitrogen) diluted in a final volume of 500ml with Opti-MEM media (ThermoFischer). To form siRNA-lipofectamine complexes, the individual solutions were incubated for 20 minutes. In this time-frame, 2.5×10^4^ cells were added in 2ml media (DMEM, 20% FBS without penicillin/streptomycin) per well in a 6-well plate. Next, 250µl of the transfection mix was added in each well. Untreated (UT) conditions were treated with 250µl of Opti-MEM medium and scrambled (sc) and siSfrp1 (si) conditions were treated with scrambled control and Sfrp1 specific siRNAs in lipofectamine solutions respectively. Furthermore, efficiency of siRNA transfection was tested over a period of 6 days, medium exchange was performed after 48 hours and then analyzed further for additional 24, 48 and 96 hours. Cells were harvested and protein isolations were carried out at each timepoint.

### Primary mouse fibroblast isolation

C57BL/6J mice were weighed first to calculate dosage of ketamine/Xylazine solution accordingly (100 μl/10 g weight). The mice were anesthetized using 1:1 ratio of ketamine/Xylazine. The mouse lungs were first flushed with 1X PBS via the right ventricle until almost white. The thorax was cleaned next and the lungs removed. The dissected lung lobes were placed in ice cold sterile 1X PBS within a 6-well plate. Using a sterile scalpel, the lobes were dissected into small 1-2mm pieces and immediately transferred into a 50ml falcon containing collagenase 1 (diluted 5mg per 50µl of 1X PBS). Collagenase digestion was carried on at 37°C at 400 rpm for 1 hour and next transferred into a 70 micron filter placed on a fresh falcon tube. Using the head of a syringe pistol the digested lung pieces were scratched onto the filter and rinsed thoroughly with sterile 1X PBS. The final suspension was centrifuged for 5 mins at 400 rpm at 4°C. Lastly, the supernatant was discarded and the pellet was resuspended in fresh cell culture medium (20% FBS and 100U/ml of penicillin/streptomycin supplemented DMEM-F12 media) and cultured under standard cell culture conditions.

### Tamoxifen preparation and treatment

C57BL/6 (B6) mice were purchased from Charles River Laboratories as already stated before. Sfrp1 Cre/ER transgenic mice were crossed with GFP transgenic mice to produce Sfrp1-GFP reporter mice. Primary fibroblasts were isolated from reporter mice as described before. 4-hydroxytamoxifen (4-OHT; Sigma Aldrich) was diluted in ethanol following the manufacturer’s recommendation. Subsequently, isolated primary fibroblasts were induced with 125nM of 4-OHT and incubated for 48 hours under standard cell culture conditions. Finally, cells were washed once with 1X PBS and fixed with 4% PFA for 30 mins at RT and imaged using a laser scanning confocal microscope (Zeiss LSM710).

### FACS sorting of Sfrp1-GFP positive cells

Using a BD Cytometry Cell Sorter FACS Aria II (Becton Dickinson, Germany), cells were suspended in MACS buffer (MACS® Separation Buffer + 2% PBS) and sorted for Sfrp1-GFP positive cells. The purity of the GFP-positive cells was subsequently evaluated using flow cytometry. For this, an aliquot of 3×10^5^ cells in 1ml MACS buffer was incubated at RT for 15 minutes with 1:500 dilution of Zombie UV™ Fixable Viability Kit (Biolegend). Cells were stained with 1 µl of anti-GFP antibody (Santa Cruz) and incubated at RT for 45 minutes. After removing the supernatant, 0.6 µl of the secondary antibody (Alexa Fluor 488; Invitrogen) was added for 30 minutes at 4°C under light-protective conditions. Prior to flow cytometry analysis, the cells were washed using MACS buffer, the supernatant was discarded, and subsequently the pellet was resuspended in 300µl of the same buffer. In live gated cells (Zombie^neg^), the FITC channel was used to assess the expression of GFP. A BD LSRII flow cytometer was utilized for flow cytometry acquisition (Becton Dickson, Germany).

### Fibroblast Invasion assay

Collagen G matrices in 96-well plates (Corning) were prepared as previously described (Oehrle et al., 2015). Briefly, succeeding gelation of the collagen, 2×10^4^ cells/well were seeded on top of the collagen plug. Cells were allowed to infiltrate the collagen matrix for 96 hours under standard cell culture conditions of 37°C and 5% CO_2_. Thereafter, each well was washed twice with PBS and subsequently fixed with 4% paraformaldehyde for 30 minutes at room temperature. Nuclei was stained with DAPI (1:1000 in PBS) overnight. The following day, z-stacks of invaded cell nuclei were imaged using a laser scanning confocal microscope (Zeiss LSM710). Furthermore, quantification of embedded cells and invasion capacity were analyzed with Imaris software (Oxford Instruments) using its spot algorithm.

### RhoA activation assay

Activation of RhoA was quantified using the RhoA G-LISA assay kit (Cytoskeleton Inc.) according to the manufacturer’s protocol. Briefly, cell lysates from cultured phFbs were prepared. Concentrations of the cell lysates were analyzed with the Precision Red™ Advanced Protein Assay Reagent (included in the kit). Absorbance was measured by a plate reader at 600 nm and a concentration range between 0.4-2.0 mg/ml was finally used. Subsequently, lysates were added to the coated wells and attachment with the Rho-GTP affinity wells were enhanced with the binding buffer. Following a series of washing steps, anti-RhoA primary antibody and subsequently secondary HRP-labeled secondary antibody were used to quantify the bound RhoA-GTP within the cell lysates. RhoA activation was finally quantified by measuring the absorbance at 490 nm using a microplate spectrophotometer (Tecan reader).

### Protein isolation and Western Blotting

Cultured cells were first washed twice with sterile 1X PBS and subsequently scraped from the wells using a cell scratcher in 200µl lysis buffer (RIPA buffer supplemented with 1x Roche complete mini protease inhibitor cocktail and Phospho-Stop phosphatase inhibitor). The cell suspension was then transferred to a vial placed on ice and further mixed on a rotor at 4°C for 1 hour. Following this mixing phase, lysates were subjected to centrifugation at 15,000 rpm for 15 minutes at 4°C to separate the supernatant containing the total protein and pellet with the cell debris. For immunoblotting, samples collected were mixed with lämli loading buffer and then separated based on molecular weight using a standard 10% SDS PAGE (20 - 30 mA per gel). Proteins were then transferred to methanol activated PVDF (Millipore) membranes (350 mA for 60 minutes). Next, membranes were blocked with 5% milk in 1xTBST (0.1% Tween®20 in TBS) and incubated with primary antibodies at 4°C overnight, followed by HRP-conjugated secondary antibodies for 2 hours at room temperature. On the day of visualization, membranes were initially washed in 1X TBST and then briefly incubated with the chemiluminescent substrates (SuperSignal®, Thermo Fisher). The following primary antibodies were used for western blotting:

### Microarray analysis

Total RNA was isolated using the PEQGold Total RNA Kit (PeqLab) according to the manufacturer’s instructions including gDNA elimination. The Agilent 2100 Bioanalyzer was used to assess RNA quality and RNA with RIN>7 was used for microarray analysis. Total RNA (150 ng) was amplified using the WT PLUS Reagent Kit (Thermo Fisher Scientific Inc., Waltham, USA). Amplified cDNA was hybridized on Human ClariomS arrays (Thermo Fisher Scientific). Staining and scanning (GeneChip Scanner 3000 7G) was done according to manufacturer’s instructions. Transcriptome Analysis Console (TAC; version 4.0.0.25; Thermo Fisher Scientific) was used for quality control and to obtain annotated normalized SST-RMA gene-level data. Statistical analyses were performed by utilizing the statistical programming environment R (R Development Core Team Ref1). Genewise testing for differential expression was done employing the paired limma *t*-test and Benjamini-Hochberg multiple testing correction (FDR < 10%). To reduce background, gene sets were filtered using DABG p-values<0.05 in at least one sample per pair and in at least two of three pairs per analysis. Array data has been submitted to the GEO database at NCBI (GSE207561).

### Fibroblast cell-morphology assessment

To prepare cells for morphology assessment, 5×10^3^ cells/well were plated in 96-well plates using DMEM-F12 media supplemented with 20%FBS and 100 U/ml penicillin-streptomycin. The cells were kept in the incubator for 24 hrs to allow cell attachment and growth under standard cell culture conditions of 37°C and 5% CO2. Subsequently, cells were fixed the next day with 4% PFA for 30 minutes at RT. Next, fixed cells were stained with DAPI (nuclear dye) and Phalloidin (label F-actin) diluted in 1X PBS and incubated overnight at 4°C. Lastly, cells were washed twice with 1X PBS and stored in fresh PBS until imaging. Afterwards, samples were washed three times with PBS for 10 minutes each. Images were acquired with a LSM 710 as z-stacks. Confocal fluorescent z-stacks were volume rendered with Imaris 7.4.0 software (Bitplane). Subsequently, the cell shape parameters were further quantified using Cell Profiler.

### Immunofluorescence and iterative stainings (4i-FFPE), microscopy, and image segmentation

For immunofluorescent staining, cells cultured in imaging-plates were fixed in 4% PFA in PBS at 37°C for 30 minutes at room temperature and then permeabilized with 0.5% Triton-X in 4% PFA for 15 minutes. Phalloidin (Thermo Fisher) (1:300) and DAPI (1:1000) were mixed in 1% bovine serum albumin (BSA, Sigma) in PBS and added to the samples and incubated at 4°C overnight. Lastly, cells were washed three times with PBS for 10 minutes each. Images were acquired with a LSM 710 as z-stacks and a LD C-Apochomat 40x/1.1 NA water objective lens (Carl Zeiss). For fibroblasts seeded on collagen gels, the cells were fixed with 4% paraformaldehyde (PFA) diluted in 1X PBS at room temperature for 30 mins. Hoechst (Pierce) was diluted in 1% BSA in PBS and incubated at 4°C overnight. Next day the cells were washed by rinsing three times with 1X PBS. Cells were finally imaged in PBS with a LSM 710 as z-stacks. Formalin-fixed paraffin-embedded (FFPE) lung tissue sections from PBS- and Bleomycin-treated mice, as well as from healthy donors and IPF patients, were first placed in a incubator at 60°C for an hour, which was followed by tissue deparaffinization process. Using a Microm HMS 740 Robot-Stainer (Thermo Fisher Scientific), the slides containing the tissue sections were automatically incubated with several chemicals as described in table S7. Next, the tissue section containing slides were placed in R-Universal buffer (Aptum Biologics) followed by transfer to an antigen retrieval buffer containing pressure chamber (2100 Retrieval, Aptum Biologics) After 30 mins inside the pressure chamber, the slides were washed once in 1X Tris buffer for 10 min and then incubated in 5% BSA in PBS for 40 mins at room temperature. Subsequently, tissue sections were stained with primary antibodies overnight at 4°C under humid conditions. Next day, the slides were washed twice in 1X PBS for 10 min, and further incubated with fluorescently-labeled secondary antibodies for 2 hours at room temperature under humid conditions. Following two additional washes, slides were then counterstained with DAPI for 1 hour at room temperature, washed again two times with 1X PBS for 10 min and subsequently dried at room temperature. Finally, tissue slides were mounted (Dako mounting medium) and kept in the dark at 4°C until further analysis.

For iterative stainings of FFPE lung tissue sections from PBS- and Bleomycin-treated mice, the iterative indirect immunofluorescence imaging (4i) procedure was adapted to FFPE sections (4i-FFPE) as originally described by Gut et. al. for cultured cells before^48^. Briefly, the FFPE sections were prepared as described above for regular immunofluorescence staining until the blocking step. Then FFPE sections were incubated in 5% BSA in PBS containing 150 mM Maleimide for 45 minutes. FFPE sections were then washed six times in 1X PBS. Next, tissue sections were stained with primary antibodies and subsequently with secondary antibodies and DAPI, similar as for regular immunofluorescence stainings. Following staining steps, slides with FFPE sections were kept in humid conditions and mounted with an imaging buffer (700 mM N-Acetyl-Cyteine in ⅘ ddH2O ⅕ HEPES. pH 7.4) with an unfixed coverslip. After imaging the coverslips and imaging buffer were gently removed, and washed 6 times with ddH_2_O. Antibodies were then eluted by incubating the samples in an elution buffer (0.5 M L-glycine, 3 M Urea, 3M Guanidium Chloride, 70 mM TCEP-HCl in ddH_2_O. pH 2.5) thrice. After each elution step samples were washed 6 times with ddH2O. Next round of staining was performed identically starting with the blocking step.

Image segmentation for the quantification of cell-type distributions over time was performed by using 1049 µm * 1049 µm (1.1 mm^2^) ROIs from whole mouse lung sections, which were stained with the appropriate antibodies for Cthrc1, Spp1, Acta2, Sfrp1, and Col28a1, and subsequently scanned with a digital slide scanner (Zeiss Axioscan 7). ROIs were cropped and imported into Imaris (Bitplane) software. During segmentation using Imaris’ surface algorithm, low intensity objects as well as objects smaller than <10 µm were filtered out. Highly large objects representing mainly structures of blood vessels and airways (e.g. smooth-muscle cells) were manually removed as objects after final segmentation. The remaining objects were counted using Imaris’ statistical tools.

### mRNA Isolation, cDNA synthesis and qRT-PCR

RNA extraction from cultured cells was performed using the PeqGold RNA kit (Peqlab) according to the manufacturer’s instructions. The concentration of the isolated RNA was assessed spectrophotometrically at a wavelength of 260 nm (NanoDrop 1000). cDNA was synthesized with the GeneAMP PCR kit (Applied Biosystems (Foster City, CA, USA)) utilizing random hexamers using 1 µg of isolated RNA for one 301 reaction. Denaturation was performed in an Eppendorf Mastercycler with the following settings: 302 303 lid=45°C, 70°C for 10 minutes and 4°C for 5 minutes. Reverse transcription was performed in an Eppendorf Mastercycler with the following settings: lid=105°C, 20°C for 10 minutes, 42°C for 60 minutes and 99°C for 5 minutes. qRT-PCR reactions were performed in triplicates with SYBR Green I Master in a LightCycler® 480II (Roche (Risch, Switzerland)) with standard conditions: 95°C for 5 min followed by 45 cycles of 95°C for 5 s (denaturation), 59°C for 5 s (annealing) and 72°C for 20 s (elongation). Target genes were normalized to HPRT expression. Primer sequences used for qPCR are detailed in Table S7.

### Code availability

No custom code was developed for the analysis. Packages and parameters used are given in the methods section.

## Supplementary Tables

Table S1: Marker gene per cell type for 10x Tcf21 PBS subset

Table S2: Marker gene per cell type for 10x Tcf21 bleomycin day14 subset

Table S3: Marker gene per cell type for Dropseq bleomycin timecourse

Table S4: Spline table for DGE genes across timepoints per cell type

Table S5: Sequences for SCRINSHOT probes

Table S6: List of antibodies

Table S7: Deparaffinization protocol

Table S8: Sequence of primers used for qRT-PCR studies

